# Control of Prion Uptake by Bone Morphogenetic Protein Signaling

**DOI:** 10.1101/2025.11.08.687161

**Authors:** Elena De Cecco, Giovanni Mariutti, Hasier Eraña, Carlos Omar Oueslati Morales, Davide Caredio, Carolina Appleton, Vangelis Bouris, Stefano Sellitto, Simone Hornemann, Carlo Scialò, Jiang-An Yin, Enric Vidal, Magdalini Polymenidou, Joaquín Castilla, Adriano Aguzzi

**Affiliations:** Institute of Neuropathology, University of Zürich, CH-8091 Zürich, Switzerland; Center for Cooperative Research in Biosciences (CIC BioGUNE), Basque Research and Technology Alliance (BRTA), Derio, Spain; Centro de Investigación Biomédica en Red de Enfermedades infecciosas (CIBERINFEC), Carlos III National Health Institute, Madrid, Spain; ATLAS Molecular Pharma S. L., Derio, Spain; Department of Quantitative Biomedicine, University of Zürich, Zürich, Switzerland; IRTA. Programa de Sanitat Animal. Centre de Recerca en Sanitat Animal (CReSA). Campus de la Universitat Autònoma de Barcelona (UAB), Bellaterra, Catalonia. Spain; IKERBASQUE, Basque Foundation for Science, Bilbao, Spain; Institute for the Science of the Aging Brain (ISAB), Lerchenfeldstr. 3, CH-9014, St. Gallen, Switzerland

## Abstract

Prions propagate through cycles of intracellular replication, release, and uptake by host cells. Each of these steps is required for disease progression. Here, we used dCAS9-VP64 mediated transactivation to screen for determinants of prion uptake. We transduced *PRNP^-/-^* SH-SY5Y cells with a post-pooled human quadruple-guide-RNA library targeting all transcription start sites of the protein-coding genome, exposed them to synthetic infectious prions, and identified modifiers of prion internalization. Specificity was tested by counterscreens with other neurodegeneration-related aggregates and endocytic probes. Surprisingly, the Bone Morphogenetic Protein (BMP) signaling pathway emerged as a strong modulator. Prion uptake was increased by transcriptional activation of the BMPR1B, BMPR2 and ACVRL1 receptors, by the BMP effector SMAD1 and by exposure to BMP ligands. Conversely, prion uptake was reduced by activation of the BMP inhibitor SMAD6. Pharmacological inhibition of BMP signaling by the antagonistic ligand noggin and by the BMP kinase inhibitor dorsomorphin decreased prion internalization, reduced the number of prion-carrying cells, and suppressed the accumulation of proteinase-resistant prion protein in a panel of chronically infected human cell lines. Hence, cell-autonomous and non-autonomous BMP signaling components regulate prion uptake, and their manipulation provides a framework for interfering with the early steps of prion propagation.

## Introduction

Prions consist of PrP^Sc^, a misfolded and aggregated version of the normal cellular prion protein (PrP^C^) which causes transmissible spongiform encephalopathies or prion diseases^1^. Prions primarily accumulate in the central nervous system, where they propagate from cell to cell along neuroanatomical pathways^2,3^ ultimately triggering neuroinflammation^4^, neuronal dysfunction^5^ and cell death. Internalization of prions by susceptible cells represents a crucial initial step in establishing iatrogenic and oral prion infections including variant Creutzfeldt-Jakob disease (vCJD^6^), and is required for the cell-to-cell spread of prions within affected organisms.

PrP^C^ is co-translationally transferred to the lumen of the endoplasmic reticulum, where it acquires a glycophosphoinositol anchor tethering it to the external surface of biological membranes. PrP^C^ conversion into prions may partly take place on the plasma membrane^7,8^, yet PrP^Sc^ is detectable in multivesicular bodies (MVBs), lysosomes and endosome-recycling vesicles^9–13^. This raises the question of how prions enter these intracellular compartments.

Moreover, prion entry receptors may play important roles in determining disease outcomes. Within the gastrointestinal tract, specialized M cells and enterocytes transport prions from the gut lumen to the gut-associated lymphoid tissue (GALT), primarily via phagocytosis and endocytosis, delivering them to follicle-associated dendritic cells where initial replication events take place^14–17^. Transepithelial transport can occur in the absence of PrP^C^ ^(14,18)^, implying the existence of additional mechanisms.

Finally, the heterogeneity of human prion diseases correlates with strain-specific patterns of cell tropism driven by variations in cellular entry routes and intracellular transport mechanisms^19^. PrP^Sc^ may propagate from infected to healthy cells through cell-cell contacts^20,21^, tunneling nanotubes^22^, exosomes^23^, and transsynaptically^24–27^. While several proteins have been proposed to act as prion receptors^28–30^, their relevance to disease is unclear and none were validated as therapeutic targets.

This landscape suggests the existence of unidentified pathways controlling prion uptake and, consequently, disease progression. How could such pathways be discovered? In most neurodegenerative diseases, the study of disease-associated genetic polymorphisms in humans has identified many modifiers of pathogenesis, yet in prion diseases this approach has proven only moderately useful^31^. Conversely, CRISPR-based genetic screens are powerful methodologies to uncover genes that modify quantitative phenotypes, such as cellular features of disease, allowing for functional interrogation and direct causal inferences^32,33^. Here, we used T.gonfio, a potent quadruple-guide (qgRNA) human CRISPR library^31^ to conduct an unbiased CRISPR-activation screen of the entire human protein-coding genome, with the aim to discover new pathways and cellular factors affecting transcellular prion spread.

## Results

### Generation of fluorescent infectious recombinant ovine prions

Infectivity is the defining hallmark of bona fide prions, and any genetic screen aimed at identifying modifiers of prion biology must rely on material demonstrably capable of transmitting prion infectivity and inducing disease. While prions purified from scrapie-affected sheep brains are highly infectious, they do not yield perfectly homogeneous preparations. Consequently, fluorescent labeling of brain-derived prions is often contaminated with non-infectious fluorescent particles. By contrast, recombinant PrP can be produced in chemically pure form, yet recombinant aggregates often fail to acquire authentic infectivity.

We therefore set out to generate prion preparations that would combine chemical purity and efficient fluorescent labeling while retaining bona fide infectivity. To this end, we produced full-length recombinant ovine PrP carrying the VRQ polymorphism, which is associated with high susceptibility to prion disease, and generated aggregates *in vitro* using Protein Misfolding Shaking Amplification (PMSA) in the presence of the cofactor dextran^34^.

Proteinase K (PK) digestion of spontaneously misfolded ovine PrP revealed the expected electrophoretic pattern for a recombinant prion with a prominent 16 kDa fragment (**Fig. 1a**). Negative staining and Transmission Electron Microscopy (TEM) of the partially purified and concentrated PMSA product showed presence of prion rods (**Fig. 1b**), with variable lengths extending up to several hundred nanometers and morphologically indistinguishable from authentic brain-derived prions. The infectivity of these aggregates was confirmed through intracerebral inoculation into TgVole (expressing bank vole PrP) and Tg338 (overexpressing sheep *Prnp*^VRQ^) mice^35,36^. All animals were euthanized upon manifestation of neurological impairment. Western blotting of PK-digested brain tissue homogenates confirmed productive prion infection and replication evidenced by the presence of PK-resistant PrP^Sc^ (**Fig. 1c**). Further inoculation of brain homogenate from a first-passage Tg338 animal to Tg338 mice induced clinical disease and PK-resistant PrP^Sc^ with electrophoretic signatures similar to the sheep scrapie isolate SSBP/1 (**Fig. 1c**). Attack rates were 100% in all bioassays with incubation periods of 198 ± 26 days post-inoculation (dpi ± SEM) in TgVole and of 367 ± 15 dpi for Tg338 at first passage. Incubation time was greatly shortened upon secondary transmission in the same model (164 ± 17 dpi) reflecting adaptation of the recombinant prion to the host (**Fig. 1d**, **Table 1**).

**Figure 1.**
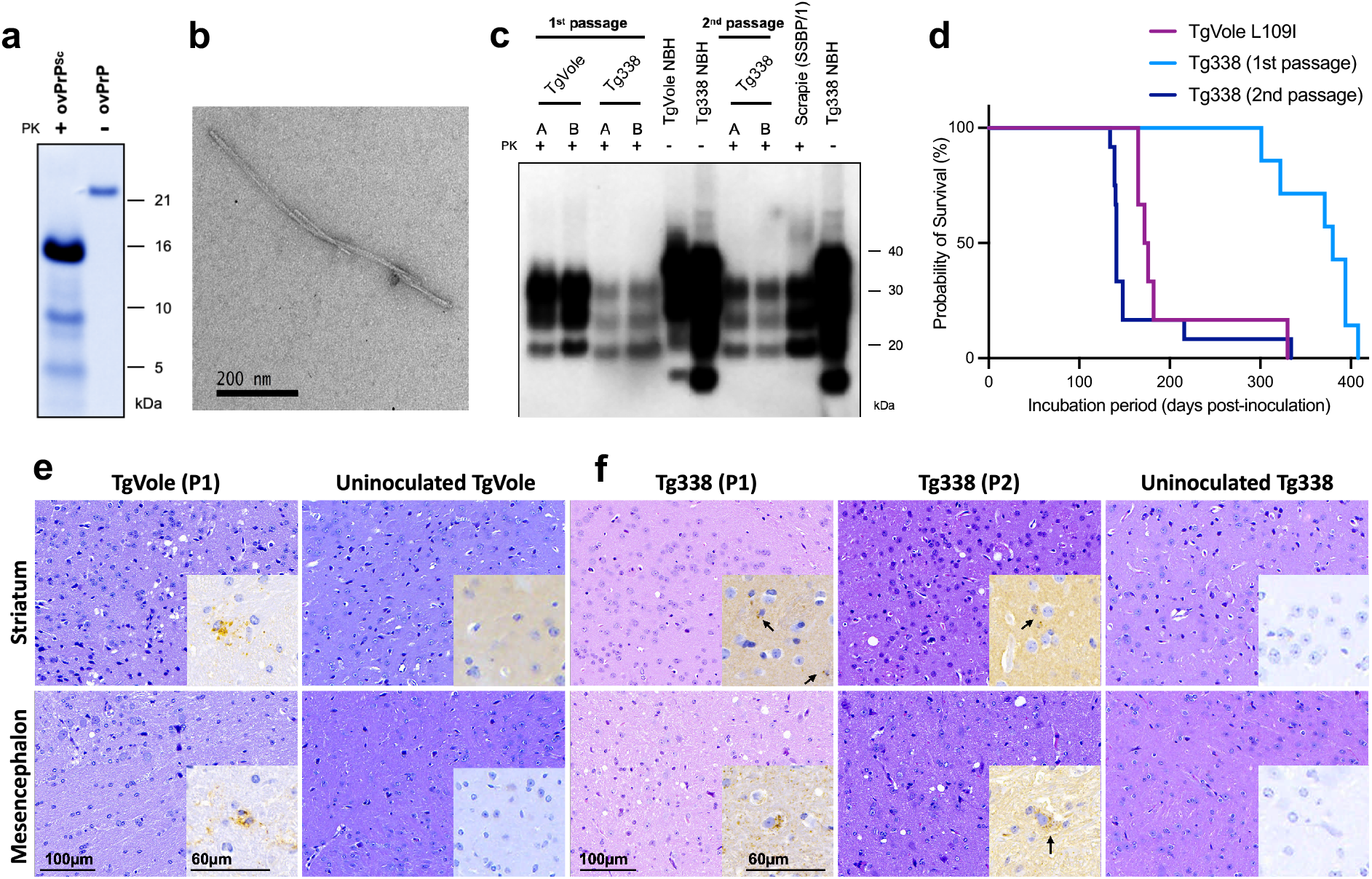
**a.** Coomassie total protein staining of PK-digested ovine recombinant PrP (VRQ variant) misfolded by PMSA, revealing a dominant ∼16 kDa PK-resistant fragment with additional ∼8–10 and ∼5 kDa bands. Monomeric ovine PrP was included as a control. **b.** TEM of partially purified PK-resistant material showing abundant fibrillar structures resembling brain-derived prion rods. **c.** Western blot of PK-digested brain homogenates from TgVole(I109)1x and Tg338 mice inoculated with PMSA-derived prions. The characteristic three-banded PrPres pattern is visible in all tested samples across both passages. PK-digested SSBP/1 and non-infectious (NBH) brain homogenates are included as controls. **d.** Kaplan–Meier analysis showing disease onset in all inoculated animals, with mean incubation periods of 198 ± 26 dpi (TgVole, n=6) and 367 ± 15 dpi (Tg338, n=7), reduced to 164 ± 17 dpi upon second passage in Tg338 (n=12). **e-f.** Histopathological staining of two brain regions (striatum and mesencephalon) from representative animals used in (d). Staining was performed using hematoxylin/eosin (nuclei and cell bodies) and anti-PrP antibodies 6C2 (for TgVole mice, panel (e)) and 2G11 (for Tg338 mice, panel (f)). Black arrows point to granular prion deposits in inoculated animals

**Table 1.** Intracerebral inoculation of Prion488 in TgVole and Tg338 mice. *One mouse died of intercurrent disease and was not included in the calculations.

| Inoculum | Mouse model | Pas-<br>sage | Attack<br>rate | Incubation<br>period<br>(dpi ± SEM) | PrP<sup>Sc</sup><br>(WB) |
| --- | --- | --- | --- | --- | --- |
| Misfolded<br>ovine PrP | TgVole(I109)1x | 1<sup>st</sup> | 6/6 | 198 ± 26 | 6/6 |
|  | Tg338 | 1<sup>st</sup> | 7/7 | 367 ± 15 | 7/7 |
| Tg338 (1st<br>passage) | Tg338 | 2<sup>nd</sup> | 12/12* | 164 ± 17 | 12/12 |

Histopathological analysis of inoculated animals confirmed that the mice had contracted prion disease with typical spongiform lesions (**Fig. 1e-f**). Immunohistochemistry using anti-PrP antibodies (6C2 for TgVole mice and 2G11 for Tg338 mice) detected faint, granular PrP^Sc^ deposits in multiple brain regions of TgVole (**Fig. 1e**) and Tg338 mice (**Fig. 1f**), most prominently in the striatum and mesencephalon. The staining intensity increased from the first to the second passage (**Fig. 1f**). Thus, ovine recombinant prions generated by PMSA are infectious and share biochemical, ultrastructural, and neuropathological characteristics with naturally occurring sheep scrapie prions. Although ovine PrP^C^ shares 89% sequence identity with human PrP^C^, there is no credible evidence that ovine prions infect humans^37,38^. The latter consideration was important for our choice of model system, as it substantially reduced the biohazards inherent to any prion manipulations.

### A PRNP-deficient human cell line expressing a programmable transactivator

The uptake of prions by susceptible cells is the initial step in the life cycle of infectious prions. We modeled this process in a clone of SH-SY5Y human neuroblastoma cells from which PrP^C^ expression was abolished by CRISPR-mediated insertion of a premature stop codon, resulting in loss of protein expression^39^. This manipulation abolishes prion replication and allows for studying the specific step of prion uptake in isolation. Importantly, removal of the human prion gene reduces any potential biohazards associated with prion replication.

For gene activation, we equipped cells with a constitutively active version of the programmable transactivator dCas9-VP64^40^ under the control of the EF1α promoter (**Fig. S1a**). To prevent any potential loss of the transgene, cells were kept under permanent blasticidin selection. *PRNP*^-/-^ dCas9-VP64-expressing cells (henceforth termed SH-KO^vp64^) showed strong genetic activation four days after transduction with quadruple CRISPR activation RNA guides (qgRNA)^31^ targeting three randomly selected genes, confirming transactivator functionality (**Fig. S1b-d**).

### Internalization kinetics of recombinant ovine prions

Prion aggregates were fluorescently labeled using AlexaFluor-488 succinimidyl ester to generate fluorescent infectious ovine prions, henceforth called prion^488^, and sonicated for 10 minutes in a high-power (∼150 W, 40 kHz) water-bath sonicator to fragment larger assemblies into smaller species. Density gradient centrifugation on 60% iodixanol showed that sonication progressively increased the prion^488^ fraction in the supernatant, indicating efficient fibril fragmentation (**Fig. S2a**). Upon flow cytometry, all aggregates showed signal in the green channel (emission ∼ 530 nm), confirming labeling (**Fig. S2c**). Sonicated prion^488^ aggregates were distinguishable from living cells by their lower forward scatter and exhibited a broad range of fluorescence intensities reflecting size heterodispersion (**Fig. S2b-c**).

We added prion^488^ (100 nM monomers equivalents) to the culture medium of SH-KO^vp64^ cells, removed adherent aggregates after 24 hours by trypsinization, and performed flow cytometry analysis. (**Fig. 2a**). We focused on prion internalization rather than surface adherence because intracellular compartments have been implicated in the prion life cycle^41^ and flow cytometry may not accurately quantify prions loosely bound to cell surfaces. Green fluorescence intensity displayed a bimodal distribution with one peak overlapping the control population (not exposed to prions) and a second peak (∼ 45% ± 4.1% of cells) with higher fluorescence. This pattern suggested that only a subset of cells internalized the aggregates. To discriminate between internalized and membrane-associated prion^488^, samples were acquired in the presence or absence of the non-cell-permeant amyloidophilic dye Trypan Blue (TB)^42,43^, which absorbs green fluorescence at 590 nm^42,43^ and suppresses signals from surface-associated prion^488^ (**Fig. 2b, Fig. S2c**). TB only modestly reduced the prion^488^-positive fraction (45% vs. 38%, p < 0.05; **Fig. 2b**), indicating that the signal primarily reflected intracellular prion^488^.

**Fig. 2.**
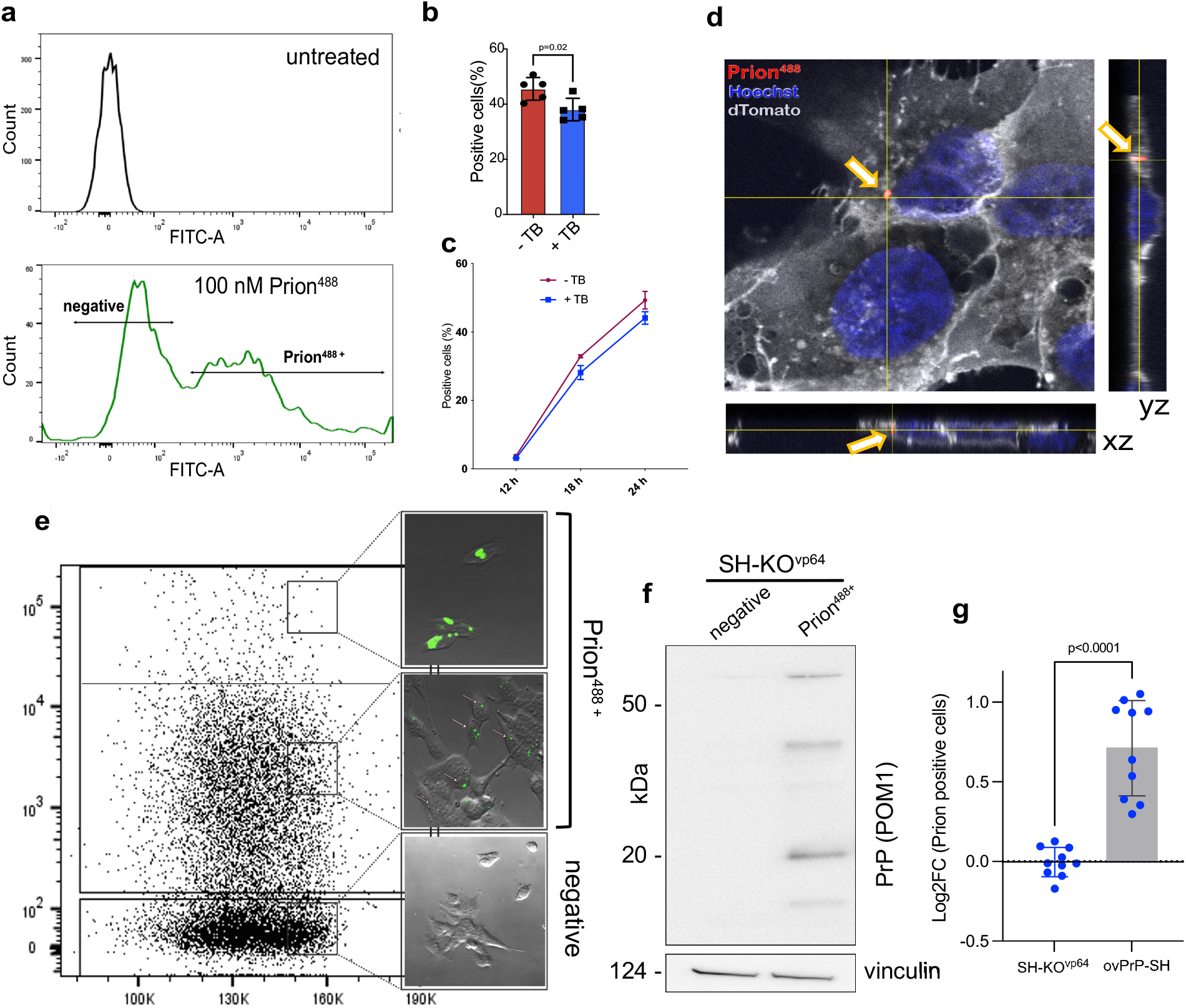
Prion uptake at single cell resolution. **a**: Flow-cytometry cell counts as a function of fluorescence intensity (FITC-A, ∼515–545 nm) for untreated (top) and Prion488-treated (bottom) cells. **b-c.** Fraction (b) and time course (c) of prion488 internalization in the presence or absence of 2mM Trypan Blue (TB). Dots: averages of two technical replicates. **d.** SH-KOvp649 cells expressing farnesylated TdTomato treated with 100 nM prion488 (24 h). Orthogonal projections of z-stack images show internalized aggregates. **e.** Prion488 treated SH-KOvp64 cells were sorted into three bins based on AlexaFluor-488 intensity, replated, and imaged. Side panels: representative images of sorted cells. **f.** Prion488+ and prion488-cells were sorted and analyzed via Western Blot. Vinculin was used as loading control. **g.** Uptake of prion488 aggregates in SH-KOvp64 and ovPrP-SH. Data are represented as log2FC calculated on the average of SH-KOvp64 cells. Dots represented individually treated wells in n=2 independent experiments. All panels: unpaired Welch’s t-test. Error bars: mean ± SD.

Consistent internalization was observed after 18 h of incubation and further increased by 24 h (**Fig. 2c**) to ≈50% of cells, which is ideal for visualizing both positive and negative modifiers of uptake. All further analyses were therefore performed at this time point. Confocal analysis of prion^488^-treated cells confirmed that aggregates were intracellular (**Fig. 2d**).

We then sorted prion^488^-exposed cells into three gated fractions with high, intermediate and no fluorescence. After sorting, re-plating and imaging, the high-and intermediate-fluorescence fractions displayed large intracellular aggregates and smaller puncta, respectively (**Fig. 2e**). This variegation of fluorescence may arise from heterogeneity in aggregate size or labeling efficiency. Alternatively, a fraction of cells may act as “super-uptakers” and internalize larger amounts of aggregates. Western Blot analysis of the prion^488-^ and prion^488+^ populations (the latter comprising pooled high-and intermediate-fluorescence fractions) confirmed enrichment of prion protein in the prion^488+^ population (**Fig. 2f**). We then conjugated prion aggregates to pHrodo Red, whose fluorescence intensity increases under acidic conditions (**Fig. S2d**). We observed a shift in fluorescence intensity in cells treated with prion^pHrodo^ (100 nM) after 24 h, but not after 1 h (**Fig. S2e-f**), consistent with our previous observations (**Fig. 2c**) and indicative of uptake and delivery of prion^pHrodo^ to acidic vesicular compartments. The fraction of pHrodo^+^ cells was comparable to that observed with prion^488^.

PrP^C^ has been implicated in the uptake of amyloids^44–50^. We expressed ovine *Prnp*^VRQ^ under the control of the EF1α promoter^51^ in SH-KO^vp64^ cells (ovPrP-SH cells; **Fig. S2g**). PrP^C^ expression resulted in increased prion^488^ uptake after 24 h treatment (**Fig. 2g**). This result is consistent with the reported role of PrP^C^ in amyloid uptake and extends it to prion aggregates, thereby providing a potential explanation for prior findings^52,53^. Finally, incubation of SH-KO^vp64^ cells with unlabelled or fluorescently labelled prion^488^ (up to 200 nM monomer equivalents) did not affect cell survival or proliferation (**Fig. S2h**). Hence, prion uptake can be robustly modelled and reliably measured in human cell lines through flow cytometry.

### Genome-wide pooled CRISPR activation screen identifies 43 new modulators of prion uptake

We then conducted a genome-wide, FACS-based pooled CRISPR activation screen in SH-KO^vp64^ cells (**Fig. 3a**). We used the *T.Gonfio* human activation library which multiplexes quadruple non-overlapping guides (qgRNAs) to enhance perturbation efficiency while minimizing off-target effects^31^. To prevent illegitimate recombination between different lentiviral transcripts, each qgRNA was individually packaged into lentiviral particles and subsequently pooled to obtain the final library. For genes with multiple transcription start site, each site was targeted individually, resulting in a pool of 22’442 lentiviral vectors targeting 19’839 protein-coding genes.

**Fig. 3.**
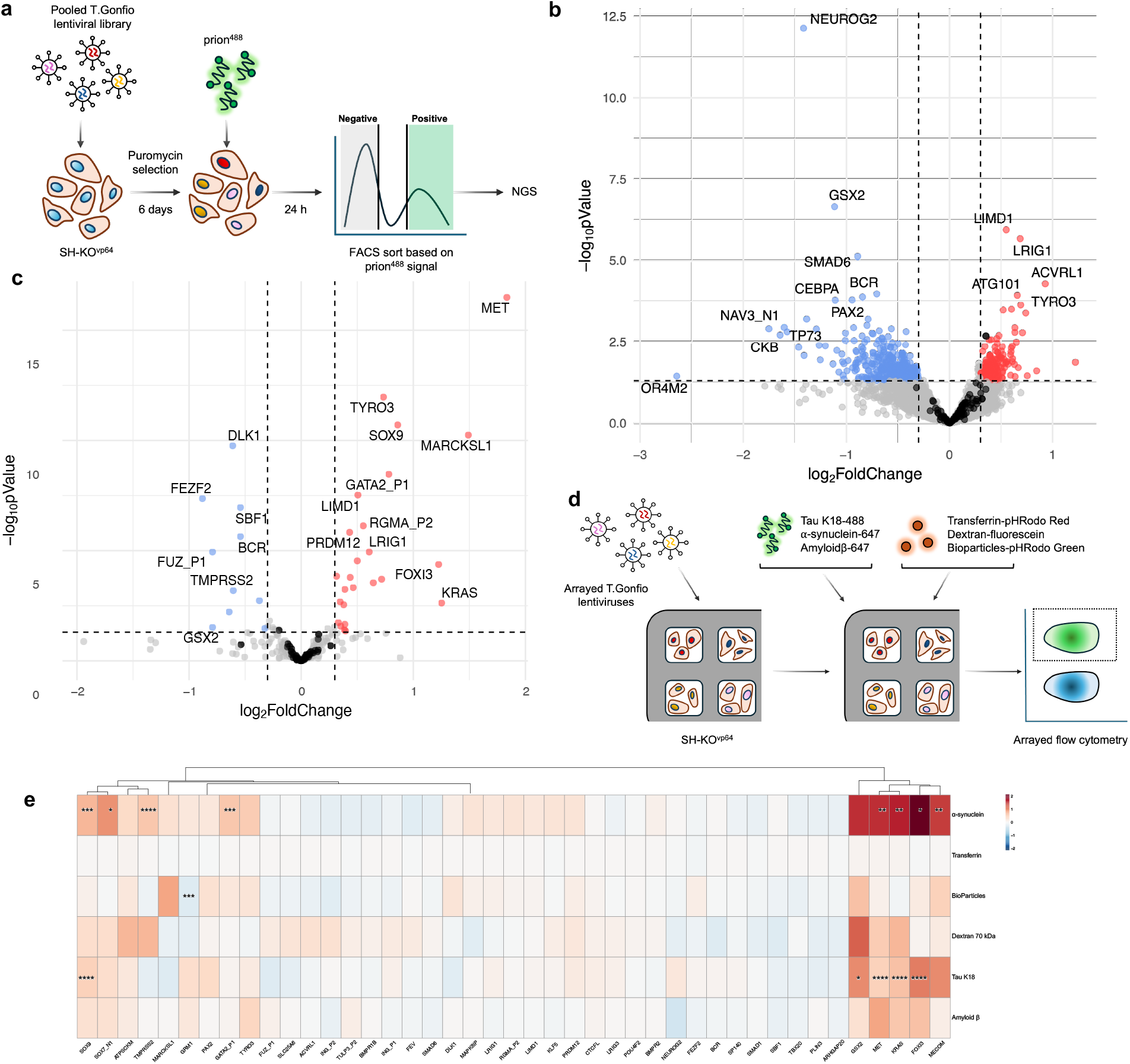
Genome-wide CRISPR activation screens for modifiers of prion uptake. **a.** Workflow of the primary screen. **b.** Volcano plot showing enriched (red) and depleted (blue) qgRNAs in prion^488+^ cells compared to prion^488-^cells. Black dots: non-targeting qgRNA. Dotted lines: thresholding criteria (|log_2_ FC|>0.3, p-value <0.05). **c.** Volcano plot showing hit genes confirmed in the secondary screen performed on the strongest modifiers. Color codes as in (b). **d.** Workflow of the arrayed CRISPRa screens on the subset of validated prion uptake modifiers. Three protein aggregates (Tau K18-488, α-synuclein-647, amyloid β-647) and three non-aggregated probes (transferrin-pHrodo Red, 70 kDa dextran-fluorescein, BioParticles-pHrodo Green) were used. **e.** Heatmap showing average log_2_FC values of three individually treated wells, calculated on MFI and normalized on the average of 10 NT controls, each in three individually treated wells. Statistical analysis: Brown-Forsythe ANOVA with Dunnett’s T3 post-hoc test. Statistics computed for genes with |log_2_FC|>0.3. *: p<0.05, **: p<0.01, ***: p<0.001, ****: p<0.0001.

SH-KO^vp64^ cells were transduced with the lentiviral library at a multiplicity of infection (MOI) of 0.3, followed by puromycin selection for five days to remove non-transduced cells. Six days after transduction, cells were reseeded, treated with prion^488^ (100 nM) for 24 hours, and sorted by fluorescence intensity at ∼530 nm) (**Fig. S3a**). Post-sort reanalysis showed that the prion^488+^ and prion^488-^populations were highly pure (**Fig. S3b**). Genomic DNA from sorted cells was extracted, amplified, and subjected to Illumina sequencing. Putative modifiers were defined by differential guide enrichment in the two sorted populations. The screen was performed twice in different days using independent cell aliquots, yielding highly correlated results (**Fig. S3c**).

Only five genes (the uptake enhancers *LRIG1* and *LIMD1* and the uptake limiters *NEUROG2, GSX2* and *SMAD6*) reached FDR-adjusted statistical significance (FDR < 0.05), but 635 genes exhibited nominal significance (p < 0.05) and meaningful effect size (absolute log_2_fold change (| log_2_FC |) > 0.3) (**Fig. 3b; Suppl. Doc. 1**). We deliberately chose a modest effect-size threshold to minimize false-negatives, while recognizing that secondary validation is required to filter potential false positives. In addition, genes with normalized read counts below 200 in both replicates of either the negative or positive population were excluded from further analysis.

Among the strongest modifiers, we found several genes involved in endocytosis, such as the TAM receptor *TYRO3*^54^ and the Ras-GTPase *KRAS*^55^, as well as genes influencing actin dynamics (i.e. *MARCKSL1*^56^). The recovery of cytoskeletal remodeling modulators is consistent with an internalization-related phenotype and supports the specificity and biological relevance of our assay. Activation of the *PRNP* gene did not increase prion uptake (log_2_FC = −0.002, p-value = 0.99), consistent with the SH-KO^vp64^ model, in which genetic ablation prevents functional PrP^C^ protein production^39^.

To refine candidate selection, we generated a CRISPR sublibrary targeting 185 of the strongest candidate genes. All alternative transcriptional start sites for these genes were included, along with 70 non-targeting guides and 52 non-hit genes as internal controls, for a total of 358 qgRNA sets. Lentiviruses were generated individually, titrated, pooled, and used to infect SH-KO^vp^^64^ cells. Screens were performed as three independent replicates on different weeks, following the same workflow as in Fig. 3a. One of the three prion^488+^ samples was excluded due to poor DNA quality. Correlation coefficients between replicates ranged from 0.96 to 0.98 (**Fig. S4a**). Out of 185 primary hits, 43 were confirmed in the confirmatory screen (|log_2_FC| > 0.3, p-value < 0.05; **Fig. 3c, Suppl**. **Doc. 2**). Importantly, only two of the 122 non-targeting and non-hit guides passed the thresholding criteria, representing false-positive rate of 0.55%.

While VP64 may activate genes adjacent to the intended qgRNA targets, chromosomal mapping of the identified hits revealed no evident topological clustering (**Fig. S4b**), suggesting that detection was unlikely to result from spurious neighboring activation.

### Prion uptake modifiers display distinct effects across endocytic cargos

Prions can conceivably enter cells through dedicated receptors, by hijacking generic cellular endocytic pathways, or by a combination of mechanisms shared with other neurodegeneration-related aggregates. To address this question, we performed arrayed CRISPR-activation screens on all validated hit genes and assessed internalization of non-prion amyloids (tau K18-488, α-synuclein-647, amyloid β-647) and three unrelated endocytic cargos (transferrin-pHrodo Red, 70 kDa dextran-fluorescein, BioParticles-pHrodo Green; **Fig. 3d**) by flow cytometry. Protein aggregates (100 nM in monomer equivalent) were incubated for 24 h, whereas non-amyloid probes were added shortly before the assay (30 min for transferrin-pHrodo Red, 2 h for 70 kDa dextran-fluorescein and BioParticles-pHrodo Green; **Fig. S5** and **Fig. S6**). Transferrin, 70 kDa dextran and BioParticles were used as markers for clathrin-mediated endocytosis, macropinocytosis and phagocytic-like endocytosis, respectively.

Unlike prion^488^ aggregates, all tested probes were readily internalized by SH-KO^vp64^ cells, resulting in >90% positive cells in cells transduced with NT control qgRNA guides (86% ± 2.6 for 70 kDa dextran-fluorescein, **Fig. S5-6**).Therefore, uptake was quantified using the median fluorescence intensity (MFI) of the positive population, which reflects the amount of internalized cargo per cell. We calculated the log_2_FC MFI values for each tested gene over the average of 10 NT controls across individual probes to compare their effect on distinct cargos and endocytic pathways (**Fig. 3e**). Sixteen genes showed moderate effect (|log_2_FC|>0.3) on at least one probe, indicating that prions exploit shared pathways to enter cells. Interestingly, none of the modifiers affected uptake of transferrin, a well-established marker for clathrin-mediated endocytosis, suggesting that this pathway is unlikely to play a major role in prion uptake. Among the protein aggregates, synuclein was the most affected by genetic upregulation of the hit genes, whereas the uptake of A β fibrils was only mildly impacted. Seven genes (*GSX2, MET, KRAS, FOXI3, MECOM, SOX9, ATPSCKM*) strongly modulated the uptake of all protein aggregates. *GSX2, MET, KRAS* and, to a lesser extent, *MECOM*, also affected the uptake of at least one non-amyloid probe, suggesting that they regulate general endocytic trafficking.

The prion uptake enhancer FOXI3, a transcription factor belonging to the family of forkhead box (FOX) proteins, strongly increased uptake of all fibrils but showed no effect on non-amyloid probes independently of their size (**Fig. 3e**) and fluorescent label. Protein-conjugated dextran uptake was unchanged, indicating that FOXI3 selectively primes cells for fibril, but not general protein, endocytosis. Hence, prion uptake may involve both shared endocytic routes and cargo-selective mechanisms.

### Modulation of BMP signaling affects prion uptake bidirectionally

Surprisingly, ClusterProfiler^57^ analysis of the primary hitlist revealed unequivocal enrichment of genes connected to the Bone Morphogenetic Protein (BMP) pathway and its downstream effectors (**Fig. S7a-b**). Core BMP components, including receptors and SMAD proteins, accounted for terms such as *“response to BMP”* and *“SMAD protein signal transduction”*. Additional morphogenetic terms (e.g., *“cartilage development”*, *“aortic valve development”*) overlapped with the BMP pathway, indicating a common signaling origin. Notably, all three LRIG (leucine-rich repeats and immunoglobulin-like domains) family members — LRIG1, LRIG2, and LRIG3 — were among the hits. Consistent with their role in promoting BMP signaling^58^, LRIG1 and LRIG3 acted as enhancers of prion uptake whereas LRIG2, whose role in BMP signaling is less well defined^58^, showed a weak inhibitory effect.

The secondary screen confirmed the enrichment of BMP-related terms observed in the primary screen (**Fig. S7c-f**). Protein-protein interaction (PPI) analysis using STRING^59^ identified two groups of interacting proteins (confidence score cutoff: 0.7), one of which comprises core members of the BMP pathway (**Fig. S7g**). When lower-confidence interactions were included, the network expanded (**Fig. S7h**), pointing to a broader transcription factor–based regulatory network that may influence cellular propensity for prion uptake. These results suggest a convergence on a single biological axis — activation of BMP/SMAD signaling— modulating prion uptake by SH-KO^vp64^ cells.

Among the cell-autonomous core components of the BMP pathway (**Fig. 4a**), three receptors (BMPR1B, log2FC: 0.46; BMPR2, log2FC: 0.71; ACVRL1, log2FC: 0.39), the BMP co-receptor RGMA (log2FC: 0.55) and the inhibitory effector SMAD6 (log2FC: −0.37) met the thresholding criteria and were confirmed as hits in the secondary screen. The positive effector SMAD1 (log2FC: 0.28) was retained for further analysis because of its well-established role in BMP signaling, making it a biologically relevant candidate. Arrayed validation was performed across multiple independent experiments using distinct prion preparations and viral stocks. All tested genes reproduced a consistent directional effect on prion uptake. As in the primary and confirmatory screens, prion uptake was increased by transcriptional activation of the receptors BMPR1B, BMPR2 and ACVRL1, as well as SMAD1, and decreased by SMAD6 (**Fig. 4b-c**). BMPR1B, SMAD1 and SMAD6 reached statistical significance whereas others exhibited more modest effects. RT-qPCR confirmed CRISPRa-mediated transcriptional upregulation of BMP pathway genes, with changes varying among targets (**Fig. S8a**). Interestingly, upregulation of noggin (NOG) reduced prion uptake in our primary CRISPR screen (log_2_FC = −0.51) (**Fig. S8g**). While pooled CRISPR screens do not generally detect paracrine factors, noggin can act in both autocrine and paracrine manners^60^. Notably, the three main repulsive guidance molecules (RGMA, which was already a confirmed hit, RGMB and HJV), which often enhance cellular BMP responses, were all positive regulators of prion uptake (RGMA, RGMB, HJV, **Fig. S8g**).

**Fig. 4.**
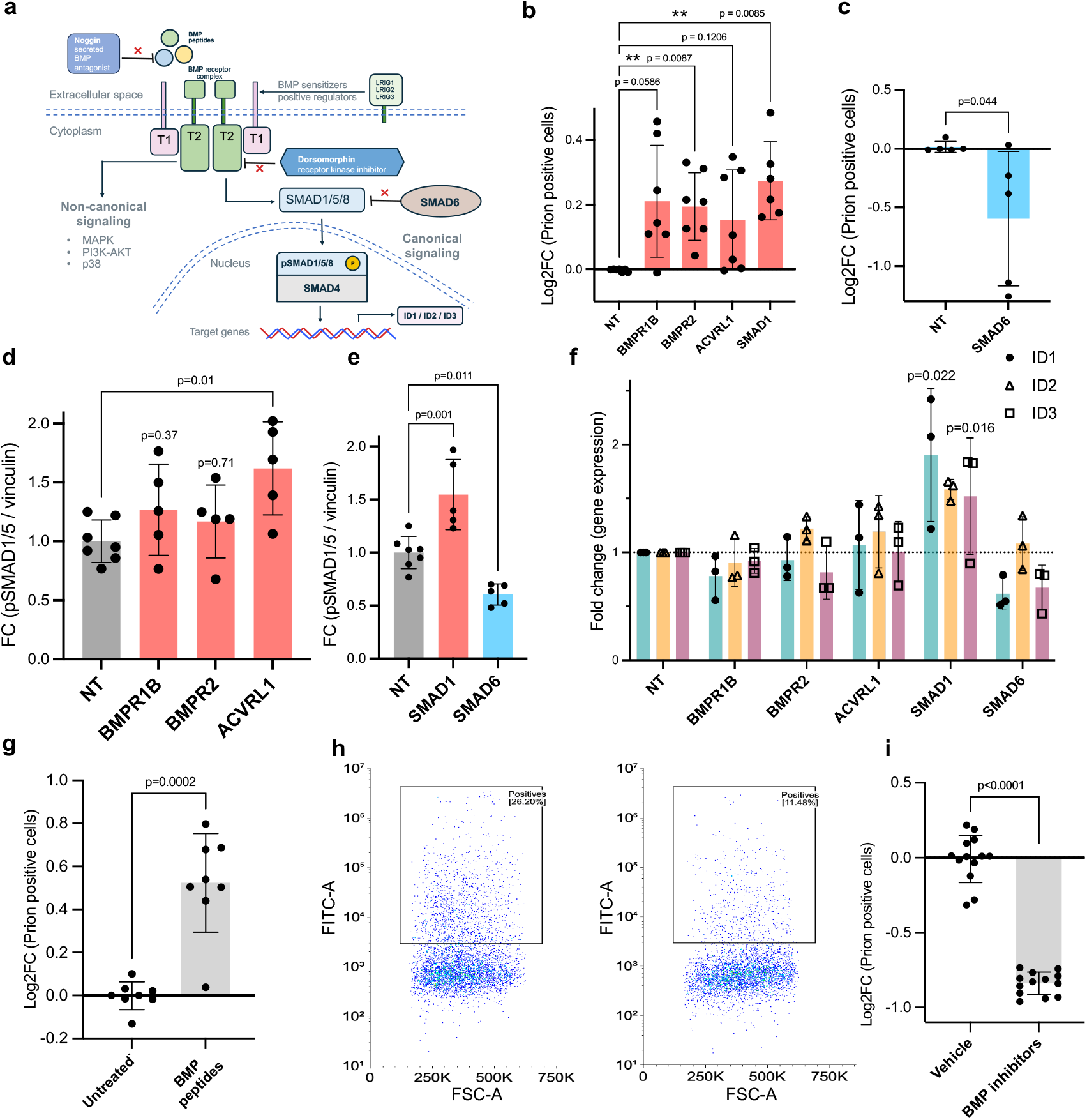
**a**. The BMP signaling pathway**. b-c** Prion^488^ uptake upon overexpression of BMP genes promoting (b) or reducing (c) prion uptake. Effect sizes: log_2_FC over non-targeting qgRNA. Dots: n=5 independent experiments (averages of 2 replicates). **d-e**. Western Blot quantification of pSMAD1/5 levels upon upregulation of BMP receptors (**d**) and SMADs (**e**). Each dot represents one genetically perturbed well. Samples were collected and analyzed on two Western blots. **f.** qPCR analysis of ID1, ID2 and ID3 gene expression upon upregulation of BMP core components. Dots: individual wells in n=3 independent experiments, each averaging 3 technical replicates. **g.** Prion^488^ uptake in SH-KO^vp64^ cells upon treatment with BMP peptides. Dots: individual wells in n=4 independent experiments, each averaging 3 replicates. **h.** Flow cytometry plots of prion^488^ uptake in SH-KO^vp64^ treated with vehicle or BMP inhibitors. **i.** Quantification of flow cytometry data shown in (h). Dots: individual wells in n=3 independent experiments. Statistical analysis: Brown-Forsythe ANOVA with Dunnett’s T3 post-hoc test (panels (b), (d9, (e), (f)); Welch’s t-test (panels (g), (i)); Student’s t-test (c). Error bars: mean ± SD.

We reasoned that overexpression of BMP pathway genes may alter downstream signaling, perhaps by increasing sensitivity to endogenous ligands. Hence, we assessed the effects of CRISPRa-mediated genetic upregulation onto the SMAD-dependent (canonical) signaling branch of BMP pathway. pSMAD1/5 levels were assessed by Western blot. Upon overexpression of the three receptors, only ACVRL1 produced modest, variable increases, whereas SMAD1 significantly increased and SMAD6 reduced pSMAD1/5, respectively (**Fig. 4d-e, Fig. S8b-c**). Next, we assessed the transcriptional activation of the SMAD-responsive ID1, ID2 and ID3 genes (**Fig. 4f**). Overexpression of BMPR1B, BMPR2, or ACVRL1 did not alter ID gene expression, whereas SMAD1 modestly increased all three, with only ID1 and ID2 reaching statistical significance. SMAD6 overexpression reduced ID1 and ID3, which was expected and consistent with partial suppression of canonical signaling. In summary, these data indicate partial enhancement of canonical BMP signaling by overexpression of BMP genes and suggest that prion uptake may also depend on non-canonical signaling pathways.

### Stimulation and repression of BMP signaling enhance and suppress prion uptake, respectively

We next assessed prion uptake following chemical modulation of the BMP pathway. We first activated BMP signaling by treating SH-SY5Y cells with a combination of three ligands (BMP2, BMP4, and GDF2). Since distinct BMP ligands can engage distinct receptor complexes and activate downstream pathways cell-state-dependently^61^, we treated cells for 4 days with a mixture of the three stimulatory ligands, to approximate the sustained pathway activation observed in the genetic screen. Under these conditions, we observed an increase in the fraction of prion-positive cells (**Fig. 4g, Fig. S8d**). In contrast, treatment with individual BMP ligands did not reproducibly enhance prion uptake (**Fig. S8e**), suggesting that the observed effect may arise from a context-dependent and combinatorial cellular response to BMP pathway stimulation.

We then suppressed endogenous BMP signaling by concomitant administration of two well-established BMP inhibitors with distinct mechanisms: noggin (a high-affinity extracellular ligand trap for BMP ligands) and dorsomorphin (which inhibits the kinase activity of BMP receptors and prevents phosphorylation of downstream targets). Both inhibitors are known to block canonical BMP signaling, and dorsomorphin has also been reported to impair non-canonical signaling (p38-MAPK)^61^. Noggin (100 ng/mL) and dorsomorphin (1 μM) were added 24 h before exposure to prion^488^ aggregates and maintained throughout the experiment. We monitored the effect of the inhibitors by quantifying pSMAD1/5 reduction via Western Blot (**Fig. S8f**). In line with our previous observations, BMP pathway inhibition led to a reduction in prion uptake (**Fig. 4i-j**). These findings are consistent with a role for BMP signaling in mediating prion internalization.

### BMP pathway inhibition reduces prion uptake and suppresses prion propagation

To test the breadth of BMP pathway-based control of prion uptake, we applied noggin (100 ng/mL) and dorsomorphin (1 μM) to the human *PRNP*-ablated glioblastoma line LN-229^62^ and evaluated prion internalization after 24 h. This treatment reduced both the fraction of prion^488+^ cells and their prion content (**Fig. 5a-c**). We then ablated *PRNP* in the human embryonic kidney-derived HEK-293A line by transient transfection of Cas9 and qgRNA targeting the human *PRNP* gene, screened individual clones for indels in the *PRNP* gene (**Fig. S9a**), and confirmed PrP^C^ absence by Western Blot (**Fig. S9b**). Next, we applied BMP inhibitors to the newly generated *PRNP*-deficient HEK-293A line and quantified prion uptake. Again, we observed a decrease in both the fraction of prion positive cells (**Fig. 5d-e**) and the MFI (**Fig. 5f**). Hence, BMP signaling plays a conserved role in modulating prion uptake across multiple human cell types.

**Figure 5.**
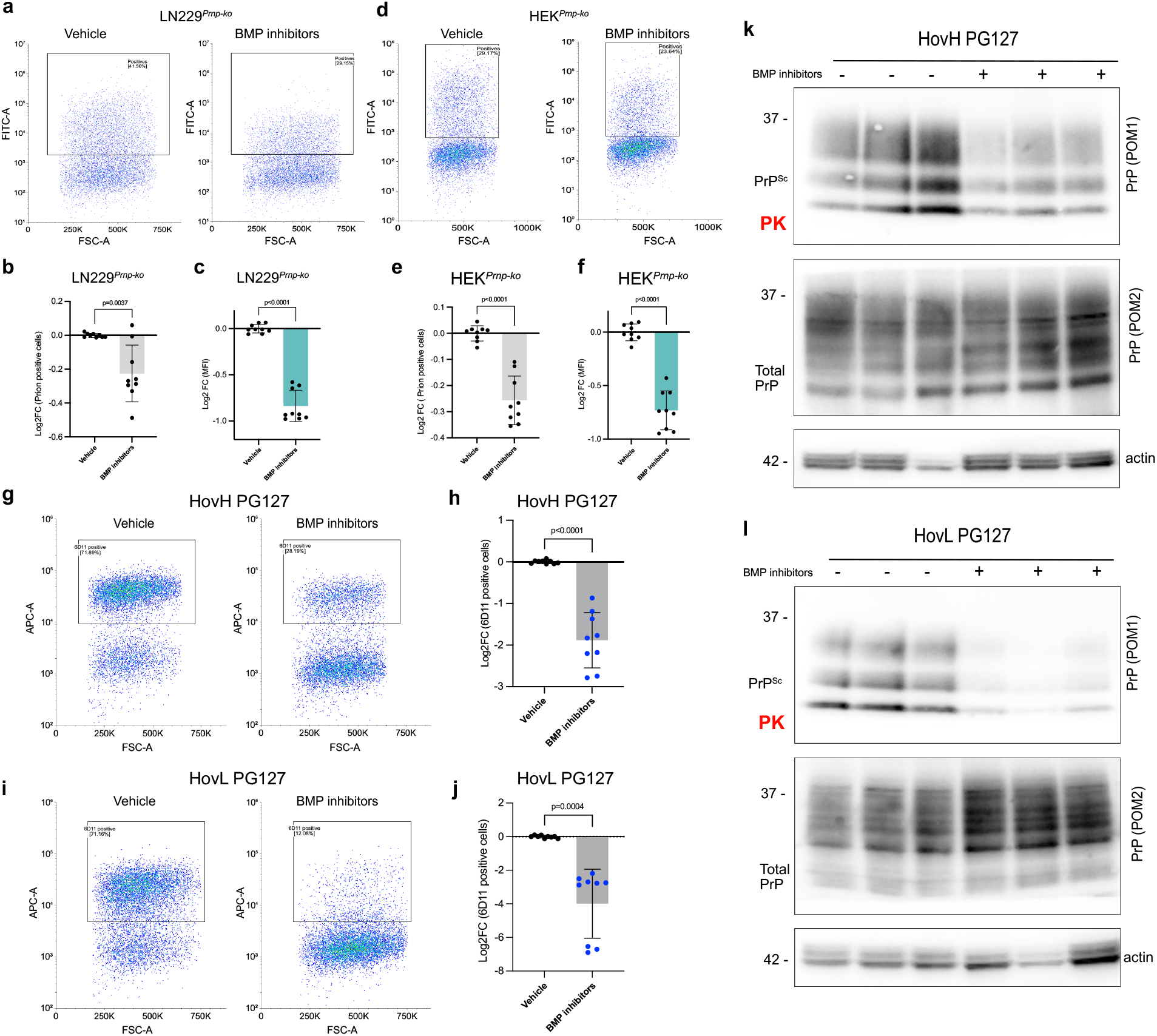
**a.** Flow cytometry of vehicle and BMP inhibitors-treated *PRNP* KO LN229 cells exposed to prion^488^ for 24 h. **b.** Prion^488^ uptake in *PRNP^-/-^* LN229 cells treated with vehicle or BMP inhibitors. Dots represent individually treated wells in n=3 independent experiments. **c.** Median fluorescence intensity (MFI) of prion^+^ cells shown in (b). **d.** Flow cytometry of vehicle and BMP inhibitors-treated *PRNP^-/-^* HEK cells exposed to prion^488^ for 24 h. **e.** Prion^488^ uptake in *PRNP^-/-^* HEK cells treated with vehicle or BMP inhibitors. Dots: individually treated wells in n=3 independent experiments. **f.** Median fluorescence intensity (MFI) of prion^+^ cells shown in (e). **g.** 6D11 staining (PrP^Sc^) of vehicle or BMP inhibited PG127-infected HovH cells. **h.** Quantification of 6D11 staining. Dots: individual wells in n=3 independent experiments. **i.** 6D11 staining (PrP^Sc^) of vehicle or BMP-inhibited HovL PG127. **j.** Quantification of 6D11 staining. Dots represent individually treated wells in n=3 independent experiments. **k.** Proteinase K (PK) digestion of samples shown in (g). Actin was acquired on a separate membrane with undigested samples. **l.** PK digestion of samples shown in (i). Actin: loading control acquired on a separate membrane with undigested samples. Statistical analysis: Welch’s t-test (panels (b),(c), (e), (f), (h), (j)). Error bars: mean ± SD.

We then overexpressed ovine PrP^C^ in *PRNP*-ablated LN-229 and HEK-293A cells (henceforth termed HovL^62^ and HovH, respectively) and exposed them to brain homogenate from mice infected (or, for control, not infected) with the ovine prion strain PG127. Both lines were found to stably propagate ovine prions over 8 passages (**Fig. S10a-b**). We then asked if pharmacological suppression of BMP signaling might impact prion levels. PG127-infected HovH and HovL cells were cultured for 14 days in the presence of BMP inhibitors (noggin 100 ng/mL, dorsomorphin 1μM). Fresh inhibitors were replenished with every medium change. Flow cytometry-based detection of PrP^Sc^ using the anti-PrP 6D11 antibody^13,63^ showed a pronounced decrease in prion infected cells upon treatment with BMP inhibitors (**Fig. 5g-h and 5j-k**) along with reduced pSMAD1/5 levels (**Fig. S10c**). PK digestion and WB confirmed the flow cytometry results (**Fig. 5i and 5l**) and indicated that both PK-sensitive and PK-resistant prions are reduced upon inhibition of the BMP pathway activity.

## Discussion

In many ways, prions behave similarly to viruses: they enter cells, co-opt host processes for replication, and spread within and between organisms. Each of these steps in the prion life cycle can be selectively interrogated with quantitative assays. Here, we set out to identify the molecular levers that control prion uptake by cells – the earliest step in prion infection. Unlike α-synuclein and tau fibrils, for which several receptors and facilitators have been described in human systems^44,45,64–69^, the molecules mediating prion engagement at the cell surface and the broader genetic networks regulating their activity are essentially unknown.

To decrease biohazards related to the use of human prions while preserving biological relevance, we employed two complementary strategies. Firstly, we used *PRNP*-deficient human neuroblastoma SH-SY5Y cells, thereby uncoupling prion internalization from downstream replication, which requires PrP^C^ expression. This enabled the study of uptake in the absence of any confounding prion amplification processes. Secondly, we generated recombinant ovine prions *in vitro* that retained infectivity in mice expressing bank-vole or ovine PrP. These aggregates recapitulated the biochemical, ultrastructural and neuropathological hallmarks of brain-derived prions^34,70^, thereby overcoming limitations associated with the impurities in brain-derived material. This approach allowed us to work with experimentally tractable yet biologically authentic prions, preserving the defining property of infectivity. Importantly, this system enables the direct interrogation of the cellular uptake of infectious prions, rather than surrogate protein aggregates, thereby providing access to an early and biologically relevant step in prion transmission that has remained difficult to study in a controlled manner. In addition, this recombinant system offers a major practical advantage, as infectious material can be generated rapidly in vitro and in large amounts, providing an essentially inexhaustible source for mechanistic studies. Crucially, transmission studies in transgenic mice did not show the emergence of human prions upon inoculation with scrapie prions^71^. These approaches provided a highly tractable low-biosafety system to interrogate prion entry.

We conjugated ovine prions with fluorophores to generate fluorescent aggregates (termed prion^488^) that could be quantitatively tracked at single-cell level. Only a subset of SH-SY5Y cells efficiently internalized prion^488^, whereas exposure to fluorescently labelled tau, α-synuclein, or amyloid-β led to more widespread uptake. This difference may reflect the relatively large size and heterogeneity of prion preparations^72^, similarly to brain-derived prions which are likely heterogeneous. While PrP^C^ is not strictly required for uptake, its expression enhanced internalization, consistent with reports implicating PrP^C^ in the uptake of diverse amyloids^44–50^. These findings extend this paradigm to infectious prions and suggest that PrP^C^ may contribute to multiple stages of the prion life cycle beyond its established role in replication and toxicity.

We performed two sequential CRISPR activations screens and identified 43 modifiers of prion uptake. These genes have not been linked to prion diseases, highlighting the ability of functional genomics to uncover pathways not readily captured by human genetics. The screens recovered multiple genes implicated in membrane trafficking, cytoskeletal organization and endocytosis, consistent with the involvement of endocytic routes. The identification of regulators of macropinocytosis (*KRAS*)^73,74^, phagocytosis (*TYRO3*)^56^ and actin-associated processes (*MARCKSL1*)^56^ among the strongest hits aligns with previous studies^75^ and indicates that a subset of prions may enter cells via phagocytosis-like pathways. Interestingly, TYRO3 is the only member of the TAM receptor family to be highly expressed in the central nervous system, and specifically in neurons^76,77^. Its overexpression was sufficient to enhance uptake, arguably through increased receptor density and activation. These results reinforce the contribution of non-canonical endocytic mechanisms to prion entry. Conversely, the enrichment in transcription factors and developmental regulators among our hits indicate that uptake efficiency may be modulated at the level of cell state. These findings suggest that susceptibility to prion uptake is not solely determined by surface-level interactions but involves coordinated transcriptional programs and broader signaling states shaping cellular permissiveness.

While some modifiers affected the uptake of multiple neurodegeneration-related aggregates (tau, α-synuclein, amyloid-β) and fluid-phase markers (high molecular weight dextran), others like the transcription factor FOXI3 displayed selectivity for fibrillar species. Clathrin-mediated endocytosis, as assessed by transferrin uptake, was largely unaffected by genetic upregulation of hit genes, suggesting that prion uptake in our system does not rely on this pathway. This implies the existence of partially distinct entry routes for proteopathic aggregates and indicates that prions preferentially exploit specific uptake mechanisms.

A central finding is the enrichment of Bone Morphogenetic Protein (BMP) signaling components among prion uptake modulators. Core pathway elements – including the receptors BMPR1B, BMPR2, ACVRL1, the co-receptor RGMA^78,79^, the effector SMAD1 and the inhibitory SMAD6, were identified in the primary screen and validated in the secondary screen and follow-up experiments. Genetic upregulation of BMP receptors or SMAD1 increased prion internalization, whereas upregulation of SMAD6 had the opposite effect. The bidirectional modulation observed across genetic and pharmacological perturbations provides strong evidence for a functional role of BMP signaling in regulating prion entry. In addition, pharmacological inhibition of BMP signaling using a combination of two well-established BMP antagonists (noggin and dorsomorphin) reduced prion uptake across multiple human cell lines of distinct origin (neuroblastoma, glioblastoma and embryonic kidney-derived lines), pointing at a more generalizable effect. While the role of BMP signaling in prion biology was not previously investigated, recent data showed that BMP4-SMAD1/5/8 signaling enhances microglial phagocytosis of amyloid-β plaques^80^. Our findings extend these observations by identifying BMP signaling as a regulator of aggregate uptake in non-microglial human cell lines and across multiple proteopathic species.

Dysfunctions in the BMP pathway have been linked to neurodegenerative diseases such as Alzheimer’s Disease (AD)^54,81,82^ and Amyotrophic Lateral Sclerosis (ALS)^83^. Although BMP signaling emerged as a prominent regulator of prion uptake in our screen (Fig. S7), we did not find evidence for enrichment of BMP pathway genes in available genetic datasets of AD (AD risk genes from a GWAS dataset, Bellenguez et al^84^), ALS (large-scale exome sequencing study, Hop et al.^85^) or Parkinson’s disease (8.9M of SNPs, Kim et al.^86^) (see Suppl. Doc. 3 for details on the analyses). The absence of a detectable signal may reflect context-dependent roles of BMP signaling or limitations inherent to cross-study comparisons, including differences in cohort composition, statistical power, variant classes, and pathway definitions. Interestingly, four of our identified modifiers of prion uptake mapped to genomic regions harboring possible risk factors for sporadic Creutzfeldt–Jakob disease (BMPR2, p-value = 0.0033; MECOM, p-value = 0.034; GSX2, p-value = 0.029; LRIG1, p-value = 0.062)^87^. Although the causal genes underlying these loci remain unresolved, this observation is consistent with the possibility that pathways influencing aggregate uptake contribute to disease susceptibility.

How does BMP signaling enhance prion uptake? SMAD1/5 phosphorylation and ID gene responses showed engagement of canonical SMAD-dependent signaling. However, BMP-dependent MAPK signaling and regulators of cytoskeletal organization could also contribute to uptake modulation. Consequently, BMP signaling may tune cellular properties—such as membrane organization or endocytic capacity—that influence aggregate uptake. In two persistently infected cell models, inhibition of BMP signaling reduced both total and proteinase K–resistant prion levels. This suggests that BMP signaling may contribute to maintaining a steady-state prion burden by modulating cell-to-cell prion transfer.

The identification of BMP signaling as a modulator of prion uptake suggests that specific cellular signaling states influence susceptibility to proteopathic seed internalization. While these findings were acquired in cultured cells, their consistency across experimental approaches suggests a broader role for host signaling pathways in shaping prion entry. These results provide a framework for the investigation of early events in prion infection and suggest that modulation of host signaling may represent a strategy to interfere with prion propagation.

## Materials and methods

### Cell culture

SH-SY5Y and SH-KO^vp64^ cell lines were cultured in OptiMEM (Gibco, 11058021) supplemented with 10% fetal bovine serum (FBS; Clontech Laboratories), 1% GlutaMAX (Gibco, 35050061), and 1% penicillin/streptomycin (P/S; Gibco, 15140122). To generate SH-KO^VP64^, a complete *PRNP*-KO SH-SY5Y clone^39^ was transduced with dCas9-VP64 (Addgene #61425). SH-KO^VP64^ cell medium was also supplemented with blasticidin 10 μg/mL (Thermo Fisher Scientific, A1113903). For lentiviral delivery in SH-SY5Y lines, cells were seeded in antibiotic-free OptiMEM and transduced in the same medium supplemented with Polybrene 8 μg/mL (Millipore, TR-1003-G). Antibiotic selection was initiated 24 hours after infection: for T.Gonfio library plasmids, puromycin 2 μg/mL (Gibco, A1113803), for farnesylated dTomato plasmid, geneticin 0.4 mg/mL (Gibco, 10131035), for dCas9-VP64, blasticidin 10 μg/mL. LN-229 and HovL cell lines were cultured in OptiMEM supplemented with 10% FBS and 1% P/S. HovL cell line medium was also supplemented with geneticin 0.4 mg/mL. HEK-293A and HovH cell lines were cultured in DMEM (Gibco, 41965039) supplemented with 10% FBS and 1% P/S. For lentiviral delivery in HEK-293A lines, cells were seeded in antibiotic-free DMEM. For HovH cell lines, the medium was supplemented with geneticin 0.4 mg/mL. HEK-293T cell line used for lentivirus production was cultured in DMEM supplemented with 10% FBS. All cells were maintained at 37 °C with 5% CO_2_ atmosphere.

### Generation of PRNP-ablated HEK and HovH cell lines

The human *PRNP*-ovinized HEK-293A (HovH) cell line was generated following the same strategy previously established for the *PRNP*-ovinized SH-SY5Y (HovS)^51^ and LN-229 (HovL)^62^ cell lines. Briefly, wild-type HEK-293A cells were co-transfected with plasmids encoding Cas9 (pSpCas9(BB)-2A-GFP; Addgene #48138) together with the CRISPRko qgRNA plasmid (from T.spiezzo library) targeting the human *PRNP* locus^31^ to disrupt endogenous expression. Clones were generated by limiting dilution, and successful PRNP deletion was verified by Western Blot, PCR amplification and sequence analysis. For the amplification of the *PRNP* gene, the following primers were used: Fwd: 5’-GCACTCATTCATTATGCAGGAAACA-3’, Rev: 5’-AGACACCACCACTAAAAGGGC-3’. *PRNP*-ablated HEK-293A cells were transduced with a lentiviral construct encoding the ovine *PRNP* gene (VRQ genotype) under the control of the EF1α promoter^51^. Clones were obtained by limiting dilution, and successful ovine PrP expression was confirmed by Western Blot. Several clones were infected with PG127-infected brain homogenate and kept in culture for 8 passages to establish a robust infection. The clone exhibiting the most reproducible and robust prion infection, along with cytoplasmic vacuolation, was selected as HovH.

### Lentivirus production

Lentiviral vectors were generated by seeding HEK-293T cells at 70% confluency and performing cotransfection 24 hours later with the cargo plasmid, along with pCMV-VSV-G (Addgene plasmid #8454) and psPAX2 (Addgene plasmid #12260), using Lipofectamine 3000 according to manufacturer’s protocol (Invitrogen). The medium was replaced 6 hours after transfection with DMEM supplemented with 10% FBS and 1% bovine serum albumin (Sigma-Aldrich, A3294). Viral supernatant was harvested 72 hours post-transfection, filtered, and stored at −80 °C. Beside the CRISPR activation library, the following plasmids were also packaged into lentivirus vectors: farnesylated dTomato plasmid, a courtesy of Dr. Nikole Zuniga Qiuroz and Dr. Esther Stoeckli^88^, dCas9-VP64 (Addgene #61425), and EF1α promoter expression vector (Sigma-Aldrich, OGS606-5U) containing the *Ovis aries PRNP* VRQ allele coding sequence (Genescript)^51^. To measure viral titer of plasmids containing fluorescent probes (i.e. CRISPRa qgRNAs), SH-SY5Y cells were seeded in 24-well plates at a final density of 100’000 cells/well. Lentiviral delivery was performed the day after seeding following the culture conditions described above. 72 hours after infection, expression of the BFP selection marker encoded by the library plasmid was measured by flow cytometry using the LSRFortessa Flow Cytometer (BD Biosciences). Data were evaluated with FlowJo 10 (Tree Star) to determine the percentage of fluorescent cells and to calculate the titer according to the following formula:

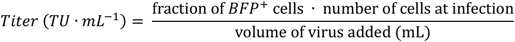

### Recombinant prion protein expression and purification

The sheep recombinant prion protein was produced based on previously published protocols^34^. The *Ovis aries PRNP* gene encoding the VRQ (amino acids 23-231) polymorphic (Accession number: AJ000738) variant was cloned into pOPIN E expression vector (Oxford Protein Production Facility) using standard molecular biology techniques. *Escherichia coli* Rosetta (DE3) competent cells (EMD Millipore) were transformed with the vector and cultured in LB medium supplemented with 50 μg/ml ampicillin at 37°C with vigorous agitation. Protein expression was induced by adding isopropyl β-D-1-thiogalactopyranoside (IPTG) to a final concentration of 1 mM when the bacterial culture reached appropriate optical density. After 3 hours of induction, the bacterial cultures were harvested by centrifugation at 4’500 g for 15 minutes at 4°C.

Bacterial pellets were resuspended in lysis buffer (50 mM Tris-HCl pH 8, 5 mM EDTA, 1% Triton X-100, 1 mM PMSF) containing 100 μg/ml lysozyme. The suspension was treated with 100 U/ml DNase and 20 mM MgCl₂ for 30 minutes at room temperature with gentle agitation. Following centrifugation at 8’500 g for 1 hour at 4°C, the inclusion bodies containing the recombinant protein were collected. These were washed with 20 mM Tris-HCl, 150 mM NaCl, 1 mM EDTA, 1% Sarkosyl, pH 8, and solubilized overnight in 20 mM Tris-HCl, 0.5 M NaCl, and 6 M guanidine hydrochloride (GdnHCl), pH 8, at 37°C with vigorous shaking.

The solubilized sheep VRQ rec-PrP was purified using histidine affinity chromatography with a HisTrap FF crude 5 ml column (GE Healthcare) connected to an ÄKTA Start FPLC system. Purification was performed by taking advantage of the natural histidine residues in the octapeptide repeat region of PrP. The column was washed with binding buffer (20 mM Tris-HCl, 500 mM NaCl, 2 M GdnHCl, 5 mM imidazole, pH 8) and the protein was eluted with elution buffer (20 mM Tris-HCl, 500 mM NaCl, 500 mM imidazole, 2 M GdnHCl, pH 8). After elution, GdnHCl concentration was increased to 6 M for long-term storage. Protein quality and purity were assessed by BlueSafe (NZYtech) staining after electrophoresis in SDS-PAGE gels. The concentration was adjusted to 25 mg/ml using 10 kDa centrifugal filter units (Amicon Ultra-15, Millipore) and the purified sheep VRQ rec-PrP was stored at −80°C until use.

### Substrate preparation for Protein Misfolding Shaking Amplification

The PMSA substrate containing sheep VRQ rec-PrP was prepared following protocols adapted from previous studies^89^. In brief, the purified rec-PrP stored in 6 M GdnHCl was thawed and diluted 1:5 in phosphate-buffered saline (PBS, HyClone) to facilitate refolding through dialysis. The diluted protein solution was dialyzed against PBS at a 1:10’000 ratio for 1 hour at room temperature to promote proper folding of the protein. Following dialysis, the sample was centrifuged at 19’000 g for 15 minutes at 4°C to remove any aggregates formed by improperly folded proteins.

The concentration of the dialyzed sheep VRQ rec-PrP in the supernatant was determined using a bicinchoninic acid (BCA) protein assay kit (Thermo Scientific) and adjusted to achieve a final concentration of 2 μM (approximately 0.1 mg/ml) in the complete substrate. The protein solution was then mixed with conversion buffer (CB: PBS containing 0.15 M NaCl and 1% Triton X-100) at a 1:9 ratio. For experiments requiring cofactor-supplemented substrate, dextran sulfate sodium salt (molecular weight 6.5-10 KDa, Sigma-Aldrich) was added to a final concentration of 0.5% (w/v).

The final substrate preparations were aliquoted into appropriate volumes to minimize freeze-thaw cycles and stored at −80°C until required for PMSA experiments. Prior to each experiment, substrate quality was verified by electrophoresis and total protein staining to ensure consistent protein concentration and the absence of degradation products.

### Coomassie staining of recombinant prion aggregates

For detection of proteinase K–resistant misfolded recombinant prion aggregates in PMSA products, 400 µL from each reaction was digested with proteinase K (PK; Roche) at a final concentration of 25 µg/mL for 1 h at 42 °C in a laboratory oven (Nahita). Following digestion, samples were centrifuged at 19’000 × g for 15 min at 4 °C, and the supernatant was carefully removed. Pellets were washed with 500 µL PBS and centrifuged again at 19’000 × g for 5 min at 4 °C. The supernatant was discarded, and pellets were resuspended in 15 µL NuPAGE LDS Sample Buffer 1× (Invitrogen).

PK-digested and non-digested samples were boiled at 100°C for 10 min and loaded onto precast gels (4–12% Bis-Tris NuPAGE gels, Invitrogen) for electrophoresis (10 min at 70 V, 10 min at 110 V, and 1 h at 150 V). Gels were subsequently stained with BlueSafe (NZYtech) for total protein detection for at least 1 h at room temperature with gentle agitation.

### Electron microscopy

Prion aggregates generated by PMSA were supplemented with SDS to a final concentration of 0.2% (w/v) and incubated for 2 h at room temperature on a rotating wheel mixer prior to processing for cryo-electron microscopy (cryo-EM). Samples were centrifuged at 200 × g for 20 min at room temperature in a swinging-bucket rotor, and the supernatant was discarded. Pellets were washed with PBS containing 0.1% SDS and centrifuged at 1’000 × g for 90 min at room temperature. Supernatant was removed, and the pellet was resuspended in 10 mM Tris–NaCl buffer with 0.1% SDS. Partially purified samples were digested with proteinase K at 60 µg/mL for 45 min at room temperature and immediately centrifuged at 1’000 × g for 30 min. The supernatant was discarded, and the pellet was resuspended in 10 mM Tris–NaCl containing 0.02% SDS.

The partially purified and concentrated sample was then loaded onto carbon-coated copper grids (Carbon Film 400 Mesh, Cu, Electron Microscopy Sciences, glow-discharged immediately before sample loading). After 1 min incubation at RT, grids were washed with deionized water for 1 min and stained with filtered 5% uranyl acetate solution for 45 s. Prion samples were then visualized in a JEM-1230 (JEOL) transmission electron microscope operated at 100 kV and equipped with an Orius SC1000 CCD camera (GATAN).

### Detection of PrP^Sc^ in inoculated animals by Western Blot

For PrP^Sc^ detection in brain homogenates from inoculated animals, proteinase K (PK; Roche) digestion was performed prior to Western blotting. Brain tissues were homogenized at 10% (w/v) in PBS (Fisher Bioreagents) supplemented with protease inhibitor cocktail (Roche) and mixed 1:1 (v/v) with digestion buffer [2% (w/v) Tween-20 (Sigma-Aldrich), 2% (v/v) NP-40 (Sigma-Aldrich), and 5% (w/v) Sarkosyl (Sigma-Aldrich) in PBS]. PK was added to a final concentration of 170 µg/mL, and samples were incubated at 42 °C for 1 h with shaking at 450 rpm in a thermomixer (Eppendorf). Digestion was stopped by addition of NuPAGE LDS Sample Buffer 4× (Invitrogen) at a 1:3 (v/v) ratio, followed by boiling at 100 °C for 10 min.

PK-digested and non-digested samples were loaded onto precast gels (4–12% Bis-Tris NuPAGE gels, Invitrogen) and subjected to electrophoresis for 1 h 20 min (10 min at 70 V, 10 min at 110 V, and 1 h at 150 V), followed by transfer onto PVDF membranes using the iBlot™ 3 system (Invitrogen). Membranes were processed using the iBind™ Flex Western device (Invitrogen) and incubated with the monoclonal anti-PrP antibody Sha31 (1:4’000; Cayman Chemical). After incubation with HRP-conjugated anti-mouse secondary antibody (m-IgGκ BP-HRP; Santa Cruz Biotechnology), signals were detected using enhanced chemiluminescence substrate (West Pico Plus, Thermo Scientific). Images were acquired with an iBright™ CL750 imaging system (Invitrogen) and processed using AlphaView software (Alpha Innotech).

### Ethics Statement

TgVole (1x) founder mice were generated at Transgenic Facility of the University of Salamanca (Spain) and breeding colonies were kept on a 12:12 light/dark cycle, receiving food and water ad libitum at CIC bioGUNE (Spain). TgVole (1x)^35^ mice inoculated in this study were obtained from the breeding colonies at CIC bioGUNE (Spain) and were inoculated at Neiker—Basque Institute for Agricultural Research and Development. This experiment adhered to the guidelines included in the Spanish law “Real Decreto 1201/2005 de 10 de Octubre” on protection of animals used for experimentation and other scientific purposes, which is based on the European Directive 86/609/EEC on Laboratory Animal Protection. The project was approved by the Ethical Committees on Animal Welfare (project code assigned by the Ethical Committee P-CBG-CBBA-0314) and performed under their supervision. The experiments involving Tg338 mice^36^, obtained from a breeding colony from at the Autonomous University of Barcelona and inoculated at the IRTA-CReSA, were approved by the animal experimentation ethics committee of the Autonomous University of Barcelona (Reference numbers: 5767 and 1124M2R) in agreement with Article 28, sections a), b), c) and d) of the “Real Decreto 214/1997 de 30 de Julio” and the European Directive 86/609/CEE and the European Council Guidelines included in the European Convention for the Protection of Vertebrate Animals used for Experimental and Other Scientific Purposes.

### Bioassays/In vivo infectivity

Spontaneously PMSA-generated, prion-containing containing sample was diluted 1:10 in sterile PBS prior to intracerebral inoculation into TgVole (1x), and Tg338 mice. For the second passage, 10 % brain homogenates from mice inoculated with the recombinant ovine prion strain was prepared as 1 % (w/v) brain homogenates in PBS from an animal selected from the first passage showing an incubation period as close as possible to the group mean.

Groups of 5 to 6-week-old TgVole (1x) and Tg338 mice were inoculated intracerebrally with 20 µl of inocula into the left cerebral hemisphere using a sterile disposable 27-gauge hypodermic needle (Terumo). Mice were anesthetized with gaseous anesthesia (Isoflurane, IsoVet®, Braun). Experimental groups were comprised of 6 to 12 animals each as indicated in the corresponding table. The animals were examined twice a week until development of neurological clinical signs, after which they were examined daily. Clinically affected animals were culled at an advanced stage of disease, but before neurological impairment compromised their welfare, by exposure to a rising concentration of carbon dioxide. The clinical signs monitored included kyphosis, gait abnormalities, altered coat state, depressed mental state, flattened back, eye discharge, hyperactivity, loss of body condition, and incontinence. Animals exhibiting two or more severe signs or invalidating motor disturbances were euthanized. Survival time was calculated as the interval between inoculation and culling or death. The brain was extracted immediately after culling and divided longitudinally; one half of the-brain was kept frozen at −80 °C for biochemical analysis and the other part was placed into 10 % phosphate-buffered formalin solution (Sigma-Aldrich) for histopathological analysis. The frozen half-brain of one of the Tg338 inoculated with the recombinant ovine prions that showed an incubation period close to the mean of the group was used to prepare the inoculum for the second passage. Briefly, the half brain was homogenized at 10% (w/v) in PBS+PI (Protease inhibitor Cocktail, Roche) and then further diluted to 1% with PBS. As for the first passage, 20 µl of the inoculum were injected intracerebrally into a new group of Tg338 mice.

### Histopathology and immunohistochemistry

Transversal sections of formalin-fixed half brains were performed at the levels of the medulla oblongata, piriform cortex, and optic chiasm for all animals inoculated intracerebrally with recombinant ovine prions and the Tg338-adapted version. All sections were then dehydrated through increasing alcohol concentrations and xylene before being embedded in paraffin-wax. Four-micrometers sections were placed on glass microscope slides and stained with hematoxylin (Sigma-Aldrich) and eosin (Casa Álvarez). Additional sections were mounted on 3-trietoxysilil-propilamine-coated glass microscope slides (DAKO) for immunohistochemistry, as previously described^34^. Treated tissue sections were deparaffinized and subjected to epitope unmasking treatments: immersion in formic acid and boiling (at pH 6.15) in a pressure cooker, followed by pre-treatment with 4 µg/ml of proteinase K (Roche). Endogenous peroxidases were blocked by immersion in a 3 % H2O2 in methanol solution. Sections were then incubated overnight with anti-PrP monoclonal antibody 6C2 (1:1000, CVI-Wageningen UR, #6C2/500) for TgVole, and with 2G11 (1:100, Bertin Pharma, #A03225) for Tg338, and visualized using the Goat anni-mouse EnVision system (DAKO) and 3,3’-diaminobenzidine (Sigma Aldrich). Images of spongiform lesions and PrPres deposits were taken from two of the most affected brain areas, striatum and mesencephalon.

### Production and labeling of synthetic recombinant ovine prions

Prion aggregates were labelled with AlexaFluor488-NHS ester (Invitrogen, cat. No. A20000) or with pHrodo Red succinimidyl ester (Thermo Fisher Scientific, cat. No. P36600) following manufacturer’s instructions to generate prion^488^ or prion^pHrodo^, respectively. Prion aggregates were centrifuged at 18’000 g at 4°C for 20 minutes to pellet aggregates. Supernatant was removed, prions were resuspended in PBS + 0.1% sarkosyl (w/v) and sonicated in a high-power water bath sonicator (Telsonic Ultrasonics) for 10 minutes (30s ON – 30s OFF cycles) at approximately 150 W and 40 kHz. For quantification, 7 mL of prion aggregates suspension were diluted in 63 uL of ultrapure GdnHCl 8M, incubated for 5 minutes at room temperature to dissociate the fibrils and then absorbance was measured at 280 nm using a benchtop spectrophotometer. To calculate the monomer equivalent concentration of each preparation, we used the following parameters: MW:22057.3 Da, molar extinction coefficient: 52495. Prion aggregates were resuspended at the final concentration of 1 mg/mL and incubated with AlexaFluor488 NHS ester (for prion^488^) or with pHrodo Red (for prion^pHrodo^) in a 1:12.5 ratio for 1 hour at room temperature on a rotating wheel, protected from light. Excess unreacted dye was removed with 4 rounds of centrifugation and resuspension in PBS + 0.1% sarkosyl. Finally, labelled prions were resuspended at the final concentration of 0.4 mg/mL (18 μM), aliquoted and stored at −20°C.

### Flow cytometry

All experiments to establish the assay for prion uptake and the genome-wide primary screen were acquired on the BD FACS Aria III (BD Biosciences). Arrayed secondary screens using non-prion aggregates and fluid-phase markers were acquired on the LSRFortessa Flow Cytometer (BD Biosciences) using the High Throughput Sampler (HTS) in Standard mode. Pooled secondary screen for validation of prion uptake modifiers was acquired on the Sony SH800 sorter (Sony Biotechnology Inc). Validation experiments of individual hits were acquired on the Sony SP6800 Flow Cytometer (Sony Biotechnology Inc) and on the Sony SH800 sorter.

For prion^488^ uptake experiments, 100’000 cells/well were plated in 24-well plates. The day after, culture medium was removed and replaced with 1 mL/well of OptiMEM + 10% FBS containing prion^488^ at 100 nM final concentration. After 24 hours of incubation, cells were washed two times with PBS, trypsinized to remove unbound prion^488^ and washed again two times in PBS before being resuspended in FACS buffer (PBS, 2% BSA, 0.5 mM EDTA, 1% Accutase) and analyzed. When checking internalization with Trypan Blue, samples were first acquired at the cytometer, then Trypan Blue was added at the final concentration of 2 mM (1:2 dilution of the commercial Trypan Blue solution, Gibco, 15250061) and samples were reacquired with the same acquisition settings.

Flow cytometry data were analyzed with FlowJo^TM^ Software (v10.4, BD Life Sciences) and with Flo-reada.io (https://floreada.io).

### Whole-genome primary CRISPR activation screen

The human T.gonfio CRISPR activation library was packaged into lentiviruses in arrayed format following the protocol described in^31^. Individual lentiviruses were pooled together, and the resulting mix was titrated as described^31^. The library was transduced into SH-KO^vp64^ at a multiplicity of infection (MOI) of 0.3 at an estimated coverage of 500 cells expressing each set of 4 guides, using transduction conditions described above. 24 hours after infection, cells were placed under selection with puromycin at 2 ug/mL final concentration to remove uninfected cells. Cells were allowed to grow for 7 days under constant puromycin selection. At 8 days from the initial seeding, cells were reseeded into culture vessels at a density of 1 cell in 0.019 mm^2^. The day after, medium was replaced with antibiotic-free OptiMEM supplemented with 10% FBS and prion^488^ at 100 nM final concentration. After 24 hours of incubation, cells were prepared for FACS sorting on the BD FACS Aria III as described above. Cells were gated using forwards and side scatter to exclude dead cells, debris and unbound prions, then gated on the area vs. height of forward scatter to isolate single cells. Singlets were then gated on the BFP signal (qgRNA library) and on AlexaFluor 488 signal (prion^488^) using qgRNA-uninfected cells and prion-untreated cells to establish thresholds. Peaks corresponding to prion^488^ negative and prion^488^ positive populations were sorted using a custom-made purity mask. Purity of the sorted populations was confirmed by subsequent reanalysis. Sorted cells were pelleted and stored at −80°C.

### Genomic DNA extraction

Cell pellets collected during primary screen were incubated overnight in Tris-EDTA pH 8 supplemented with SDS (2% final concentration) and proteinase K (2 mg/mL final concentration) at 50°C and 500 rpm constant shaking. The next days, samples were transferred to new tubes, two volumes of buffer containing chaotropic salts (6M Gdn-SCN, 50 mM sodium citrate, 1M Tris, pH 8) (for 10^7^ cells, 2.5 mL of lysis buffer and 5 mL of buffer were used) were added, and samples were incubated at 37°C under slow constant stirring for at least 2 hours. These two steps ensure a thorough cell lysis and the removal of prion infectivity. Next, samples were subjected to three subsequent rounds of extraction using phenol:chloroform:isoamyl alcohol (25:24:1) consisting of 30 min incubation with 1.5 volumes of Ph:Chl:IAA and 30 min of centrifugation at 5’000 g to separate phases. After each centrifugation, the aqueous phase containing gDNA was transferred to clean tubes, paying attention to not disturb the interphase. To remove residual phenol, a fourth extraction step consisting of a 30 min incubation step with chloroform followed by centrifugation for 10 min at 5’000 g was performed. The recovered phase was then mixed with cold ethanol (2 volumes), 4% (v/v) sodium acetate (Sigma) and GlycoBlue (Invitrogen), and incubated overnight at −20°C. Next, gDNA was precipitated by centrifugation at 5’000 g for 30 min. gDNA pellets were washed three times in excess 70% ethanol (15 min incubation followed by centrifugation for 10 min at 10’000g), then left to dry to remove residual ethanol and resuspended in 10 mM Tris-HCl pH 8. To ensure proper resuspension, samples were incubated in buffer overnight at 37°C before being collected. gDNA quantification was performed using Qubit (ThermoFisher Scientific, 1X dsDNA Broad Range Quantification Kit, Q32853).

### qgRNA guides and NGS sequencing

qgRNAs were amplified by PCR reactions as previously described^31^. The following barcoded forward primers were used:

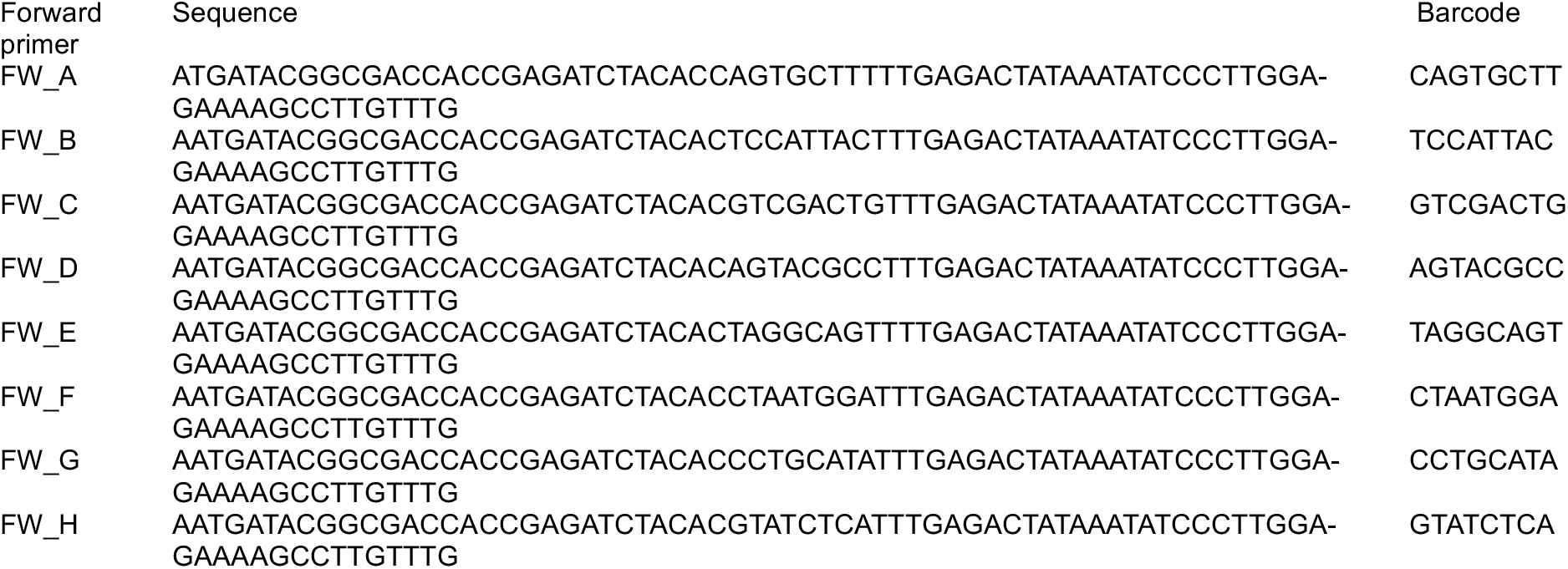

Clean-up of PCR products from reaction components and primer dimers was performed using SPRIselect Beads (Beckman Coulter, B23317) following manufacturer’s instructions. Samples were pooled together and sequenced on an Illumina NextSeq 2000 instrument using a P2 flowcells and a 300 cycles Illumina Cartridge.

### Secondary screen

Selected candidate genes were manually picked from the T.gonfio arrayed library in the form of glycostocks, amplified^31^, and distributed in four 96 deep-well plates. For plasmid extraction, 50 μL of bacterial cultures were transferred in new 96 deep-well plates containing 1.2 mL of Terrific Broth with 15 µg/ml trimethoprim (Sigma, cat. #T7883-5G) and 15 µg/ml tetracycline (Sigma, cat. #T3258-5G) using ViaFlo (Integra). Plates were grown for 48 h at 30°C and 1’100 rpm shaking on a thermal shaker (http://www.rooyn.com/products/446/2459.aspx, #RS300). Bacteria were pelleted by centrifugation at 4’000 rpm for 20 min and pellets were resuspended in 200 μL P1 buffer followed by 200 μL of lysis buffer P2 and 200 μL of neutralizing buffer P3. After each step, samples were mixed using ViaFlo to ensure proper reagent mixing. Plates were then centrifuged at maximum speed for 10 minutes to pellet chromosomal DNA and bacterial debris. Supernatants were transferred to new 96 deep-well plates and 2 volumes of cold ethanol were added. Plasmid DNA was precipitated by centrifugation at maximum speed for 10 min, then pellets were air-dried and resuspended in 50 μL of ddH_2_O by vortexing. DNA was bound to Sera Mag Select Beads by incubation at room temperature for 10 minutes and washed for two times with 1 mL of 70% ethanol. Beads were then air-dried and bound DNA was eluted in 125 μL of Tris-EDTA pH 8. To ensure proper elution, plates were incubated at 65°C for 10 minutes before transferring the eluted DNA to new plates. 10 randomly selected wells per plate were quantified using Nanodrop. The resulting averaged concentration was applied to the entire plate.

Individual plasmids were packaged into lentiviruses in arrayed format as done for the genome-wide library. A portion of the lentiviral-containing supernatants (30 mL/lentivirus) were pooled to generate the secondary screen pooled library, while the remaining volume was aliquoted into arrayed 384-well plates and stored at −80°C.

Pooled secondary screen was carried out following the same protocol of the genome-wide primary screen, except for the starting coverage that was expanded to 5000X. Differently from the genome-wide screen, FACS sorting of the secondary screen samples was carried out on a Sony SH800 sorter (Sony Biotechnology Inc.). The screen was performed in three independent replicates and samples were sequenced all together on an Illumina NextSeq 2000 instrument (Illumina).

### Cytotoxicity assay

Cells were seeded in 96-well plates with opaque wells at the final density of 4’500 cells/well. The day after, cells were treated with increasing concentrations of unlabelled prion aggregates or prion^488^ for 24 hours. Viability was measured using CellTiter-Glo 2.0 (Promega) and read on the Envision plate reader (Revvity).

### Purification of recombinant neurodegeneration-related proteins (TauK18, α-synuclein, Amyloid-β 1-42) and preparation of pre-formed fibrils

Monomeric TauK18 fragment was purified as described previously^44^. Aggregation was carried out in a FLUOstar Omega (BMG Labtech) plate reader following the aforementioned protocol ^44^. Monomeric α-synuclein was purified as described in Neupane et al.^90^. Aggregation was carried out in a benchtop thermoblock following a published protocol^91^. Aβ_1-42_ peptide was purified following a modified protocol as described^92,93^. Aggregation was carried out in a benchtop thermoblock following the same protocol^92,93^.

### Arrayed secondary screens for endocytosis modifiers

SH-KO^vp64^ cells were seeded in 96-well plate at a density of 10’000 cells/well. The day after, cells were transduced with arrayed lentiviruses at an MOI of 3. Transduction was carried out in OptiMEM + 10% FBS + Polybrene 8 μg/mL, and a reduced volume of 40 μL/well was used. One day after transduction, virus-containing medium was replaced with fully supplemented OptiMEM. Cells were cultured for 5 days after transduction and then treated with either endocytic probes or amyloid aggregates. Treatments were performed as follows:

| Treatment | Final concentra- | Incubation time (h) |
| --- | --- | --- |
| Tau K18 PFFs-488 | 100 nM | 24 |
| α-synuclein PFFs-647 | 100 nM | 24 |
| Amyloid b-647 | 100 nM | 24 |
| Dextran-Fluorescein 70 kDa | 125 μg/mL | 2 |
| Transferrin-647 | 500 nM | 0.5 |
| *E.coli* pHRodo Green Bioparticles | 100 μg/mL | 2 |

The day of the analysis, cells were washed 2 times with PBS, then detached with Trypsin and neutralized with FACS buffer containing 2% FBS/PBS. After thorough in-well mixing, cells were analyzed using a LSRFortessa Flow Cytometer (BD Biosciences) equipped with a High Throughput Sampler.

### Immunoblotting

Cell lysates were prepared in lysis buffer (50 mM Tris–HCl, pH 8, 150 mM NaCl, 0.5% sodium deoxycholate, and 0.5% Triton-X 100) supplemented with protease inhibitors (cOmplete Protease Inhibitor Cocktail, Sigma-Aldrich). The bicinchoninic acid assay (BCA) was used to measure the total protein concentrations according to the manufacturer’s instructions (Pierce). Samples were boiled at 95°C in NuPAGE LDS Sample Buffer 1x (Invitrogen) supplemented with 1 mM DTT (Sigma-Aldrich), loaded onto precast gels (4-12% Bis-Tris NuPage gels, Invitrogen), and blotted onto PVDF membranes (Invitrogen).

For PK-digested samples, 30 μg of total protein lysate was diluted lysis buffer without protease inhibitors to 20 μL final volume. 2 μL of PK (stock concentration 25 μg/mL) were added to the samples. Samples were incubated in a thermoblock at 37°C with 700 rpm shaking for 30 min. Digestion reaction was stopped by addition of 10 μL NuPAGE LDS Sample Buffer 4X containing 20 mM DTT (final DTT concentration in the sample: 5 mM) and boiling at 95°C for 10 min. For undigested samples, 15 μg of total lysates were mixed with NuPAGE LDS Sample Buffer 4X containing 20 mM DTT and boiled at 95°C for 10 min.

The following antibodies were used: anti-PrP Sha31 1:4000 (Bertin Pharmaceuticals, #A03213), anti-Cas9 1:1000 (Cell Signaling, #14697), anti-actin HRP conjugated 1:10’000 (Sigma-Aldrich, A3854), anti-Rab13 1:1000 (Abcam, ab180936), anti-ADAM10 1:1000 (Abcam, ab124695), anti-vinculin 1:1000 (Abcam, ab129002), anti-PrP POM2 0.3 mg/mL, anti-PrP POM1 0.3 mg/mL, anti-phospho SMAD1/5 (CellSignaling #9516), anti-Mouse-HRP 1:10’000 (Jackson ImmunoResearch #115-035-005), anti-rabbit-HRP 1:10’000 (Jackson ImmunoResearch #705-035-147), anti-CD230 (6D11)-647 antibody (BioLegend, #808008).

### Treatment with BMP peptides

BMP2 (Gibco, cat. No. 120-02-10UG), BMP4 (Gibco, cat. No. PHC9534-10UG) and GDF-2 (Gibco, cat. No. 120-07-10UG) were purchased in lyophilized form, resuspended in 0.1% BSA/PBS at 100 μg/mL and stored at −20°C. For cell treatments, BMP peptides were diluted into culture medium at the final concentration of 33 nM. When added together, each BMP peptide was present in the culture medium at 33 nM final concentration. Peptides were supplied daily. Every two days, exhaust medium was replaced by fresh culture medium.

### Treatment with BMP inhibitors

Noggin (Fisher Scientific, cat. No. 6057NG025CF) was resuspended in 0.1% BSA/PBS at the final concentration of 250 μg/mL and stored at −20°C. Dorsomorphin (Sigma Aldrich, cat. No. P5499-5MG) was resuspended in DMSO at the final concentration of 5 mM and stored at 4°C. For cell treatments, inhibitors were diluted into culture medium at the final concentration of 100 ng/mL (noggin) and 1 μM (dorsomorphin). BMP inhibitors (PBS and DMSO in the vehicle-treated samples) were supplied every two days in fresh culture medium.

### PrP^Sc^ detection by 6D11 antibody staining

Prion-infected cells were dissociated and fixed with Cyto-Fast^TM^ Fix/Perm buffer set (BioLegend, 426803) for 30 minutes at RT. Samples were then incubated with 3.5 M guanidinium thiocyanate for 10 minutes, washed immediately with Cyto-Fast^TM^ Perm/Wash solution and stained with AlexaFluor647-conjugated mouse monoclonal antibody 6D11 (BioLegend, 808008) diluted 1:200 in 1X Cyto-Fast^TM^ Perm/Wash solution. Samples were acquired on a Sony SH800 sorter (Sony Biotechnology Inc).

### Statistical Analysis

#### CRISPR screens

Data analysis was performed using the Sushi platform^94^ (FGCZ, University of Zurich) and R (version 4.5.1). Reads quality was assessed using FastQC tool (version 0.12.0)^95^. One sample from the secondary screen with abnormally high GC% (>70%) content was excluded from further analysis. qgRNA-2 and qgRNA-3 sequences were mapped to the Human T.Gonfio CRISPR activation library to quantify their absolute abundance in each condition and to identify their corresponding target genes. Reads were normalized across conditions using the median ratios method implemented by the R package DESeq2 (version 3.22)^96^. Differential guide enrichment was performed using DESeq2^97^. Candidate hits were defined according to the following criteria: p-value (unadjusted) ≤ 0.05, and |log_2_FC|≥0.3 for downregulators and upregulators, respectively. Correlation between replicates was expressed as Pearson’s Coefficient (r). Over Representation Analysis (ORA) on candidate hits was performed using the R package ClusterProfiler (version 3.2)^57^. The genomic localization of hits was visualized with a custom script developed by A.A. (https://github.com/aagaag/ManhattanCrispr).

Supplementary Fig. 8G – Limma moderated interpretation of screen results: each sgRNA was treated as an independent test and two replicates (positive and negative) were used to estimate both effect size and uncertainty. For each sgRNA, raw counts from prion-positive and prion-negative wells were converted into replicate log_2_ fold changes and effect size was defined as the mean of the two replicates.

With only two observations per sgRNA, the variance estimate is unstable, and raw t-tests may give inflated significance for targets whose replicate values happen to differ more than average. To address this, we used the limma (Linear Models for Microarray Data) empirical Bayes framework^98^ in which the log-transformed variance of each sgRNA (rep1 - rep2)^2^ / 2 is adjusted for the expected bias of a chi-squared-based variance estimator. The distribution of these corrected log-variances across all sgRNAs is used to estimate a global prior variance and prior degrees of freedom. The prior parameters summarize the typical scale and spread of variance estimates in the dataset. Each sgRNA’s variance is then shrunk toward this prior by a weighted average, producing a posterior variance that is less influenced by noise in any single replicate pair. The moderated t-statistic is then calculated as the mean effect divided by the moderated standard error, and the resulting two-sided p-value is obtained using t-distribution degrees of freedom augmented by the prior. It is preferred over raw p-values because it balances per-sgRNA uncertainty with global variance behavior and reduces the impact of extreme noise in individual cases. The volcano plot uses mean log2 fold change on the x-axis and ‘-log10(p_value_limma_moderated)’ on the y-axis.

#### Individual experiments

statistical significance was evaluated using the following statistical tests: 1) Brown-Forsythe ANOVA with Dunnett’s T3 post-hoc test (Fig. 3e, Fig. 4b, Fig. 4d-f, Fig. S3h, Fig. S8e); 2) two-sided Student’s t-test (Fig. 2b, Fig. 4c); 3) two-sided Welch’s t-test (Fig. 2g, Fig. 4g, Fig. 4i, Fig. 5b-c, Fig. 5e-f, Fig. 5h, Fig. 5j). The number of replicates is indicated in the figure legends.

### Test statistics

Fig. 2b: t=2.892, df=8

Fig. 2g: t=7.235, df=10.68

Fig. 3e: TauK18: F*=18.73, DFn=44.00, DFd=29.91; α-synuclein: F*=44.65, DFn=44.00, DFd=35.98; amyloid-β: F*=8.324, DFn=44.00, DFd=54.73; dextran-fluorescein: F*=2.97, DFn=44.00, DFd=54.26; BioParticles: F*=11.56, DFn=44.00, DFd=48.80; Trasferrin: F*=8.439, DFn=44.00, DFd=48.72

Fig. 4b: F*=4.565, DFn=4.00, DFd=20.57

Fig. 4c: t=2.382, df=8

Fig. 4d: F=0.9719, DFn=3, DFd=18

Fig. 4e: F=2.028, DFn=2, DFd=14

Fig. 4f: ID1: F=1.195, DFn=5, DFd=12; ID2: F=0.6476, DFn=5, DFd=12; ID3: F=0.4606, DFn=5, DFd=12

Fig. 4g: t=6.228, df=8.104

Fig. 4i: t=17.09, df=17.16

Fig. 5b: t=4.025, df=8.082

Fig. 5c: t=14.24, df=9.186

Fig. 5e: t=7.892, df=9.569

Fig. 5f: t=11.05, df=10.89

Fig. 5h: t=8.471, df=8.063

Fig. 5j: t=5.862, df=8.024

Fig. S3h: F*=0.6034, DFn=7, DFd=12.21

Fig. S8e: F*=2.020, DFn=3, DFd=3.764

## Authors’ contributions

E.D.C.: designed, performed, contributed to, supervised and coordinated all experiments and analyses; execution of primary and secondary screens; data curation and visualization; manuscript writing. G.M.: performed bioinformatic analysis, data curation and visualization of primary and secondary screens, except for Fig. S8G; contributed to all experiments shown in Fig. 1 to Fig. 5; assisted in experimental design and manuscript writing. C.M.: generated the HovH and HovL cell models; performed prion infection and maintenance of all prion-infected cell lines. D.C.: performed bioinformatic analysis, data curation and visualization of the primary screen (Fig. 3, Fig. S4), except for Fig. S8G. H.E.: purified recombinant ovine PrP; produced synthetic prionaggregates and performed their biochemical characterization, including Transmission Electron Microscopy imaging (Fig. 1). C.A.: executed and assisted with validation experiments (Fig. 3-4). V.B.: assisted with experiments shown in Fig. 6. S.S.: data curation and visualization, manuscript writing. S.H.: assisted with the execution of the secondary screen (Fig. 3); purified Aβ peptide (Fig. 3); manuscript writing. C.S.: generated the farnesylated dTomato cell line used in Fig. 1F. J-A.Y.: designed and produced the T.gonfio CRISPR activation library. E.V.: inoculated mice with synthetic ovine prions and carried out all in vivo experiments to determine infectivity (Table 1). M.P.: manuscript writing and correction. J.C.: produced synthetic prion aggregates; data curation and visualization; manuscript writing. A.A.: project design and supervision; secured funding; generated the Manhattan plot in Fig. S5 and several bioinformatic tools for quality control; performed comparative genomic analyses with AD, ALS and PD datasets; performed the statistical analysis on the primary screen reported in Fig. S8G; wrote and debugged the scripts for the generation of Fig. S4B (Appenzeller plot); wrote parts of, and corrected, the manuscript.

## Funding and acknowledgements

E.D.C. was recipient of the UZH postdoctoral grant (2020), the FreeNovation grant (2021-2022), the Synapsis Career Development Award (2022-2024, CDA-01), and is co-applicant with A.A. of the Parkinson Schweiz Grant (2023), the SNSF Projects and Applications Grant (2024) and the Uniscientia Grant (2025). J-AY received a UZH postdoctoral grant (2021) and a Synapsis Career Development Award. S.H. was supported by the Michael J. Fox Foundation (MJFF-020710, MJFF-021073) and is supported by the Foundation for Research in Science and the Humanities at the University of Zurich (STWF-24-11) and the Innovation Fund of the University Hospital of Zurich (INOV00169). A.A. was supported by a Distinguished Scientist Award of the NOMIS Foundation and grants from the Michael J. Fox Foundation (grant ID MJFF-022156 and grant ID MJFF-024255), the Swiss National Science Foundation (SNSF grant ID 179040 and grant ID 207872, Sinergia grant ID 183563), the GELU Foundation, swissuniversities (CRISPR4ALL), the Human Frontiers Science Program (grant ID RGP0001/2022), the Parkinson Schweiz Foundation and the Uniscientia Foundation. We thank the following for their support: IKERBasque foundation, vivarium and maintenance from CIC bioGUNE and the personnel from (IRTA-CReSA), especially Dr. Enric Vidal and Samanta Giler, and the personnel from NEIKER for technical support. We thank Dr. Edoardo Marcora for help on data analysis, Dr. Simon Mead for sharing the list of sCJD-associated genomic loci99, Dr. Ilan Margalith, Joanna Zukowska and Sebastian Hachenberg for their help with lentivirus production, Andrea Armani and Geoffrey Howe for discussions, as well as Dr. Susanne Kreutzer, Dr. Adriana Hotz and the Functional Genomics Center Zurich (FGCZ) for preparing sequencing libraries, CRISPR screen NGS sequencing, RNA Sequencing, quality control and technical support. The funders of the study had no role in study design, data collection, data analysis, data interpretation and manuscript writing. The corresponding authors had full access to all the data in the study and had final responsibility for the decision to submit for publication. Statistical analysis and graphs were generated with R (version 4.5.1) and GraphPad Prism version (10.2.0, GraphPad Software).

---

**Supplementary Figure 1.**
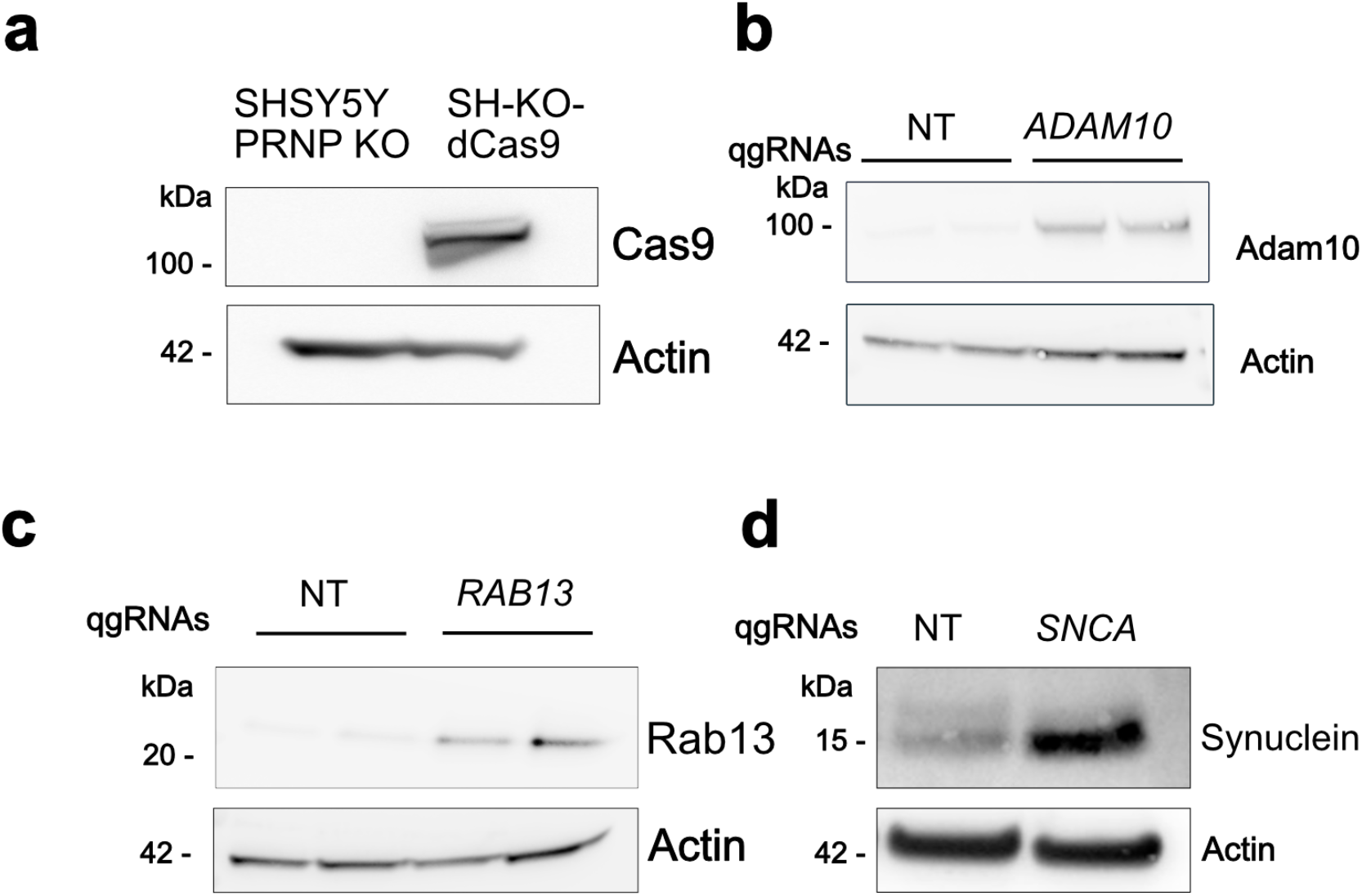
**a.** Western Blot shows expression of the transactivator dCas9-VP64 in transduced cells compared to control cells. **b-d.** Overexpression of Adam10 (**b**), Rab13 (**c**) and α-synuclein (**d**) after 4 days from qgRNA lentiviral transduction confirms activity of the transactivator dCas9-VP64. Actin was used as loading control in all Western Blots.

**Supplementary Figure 2.**
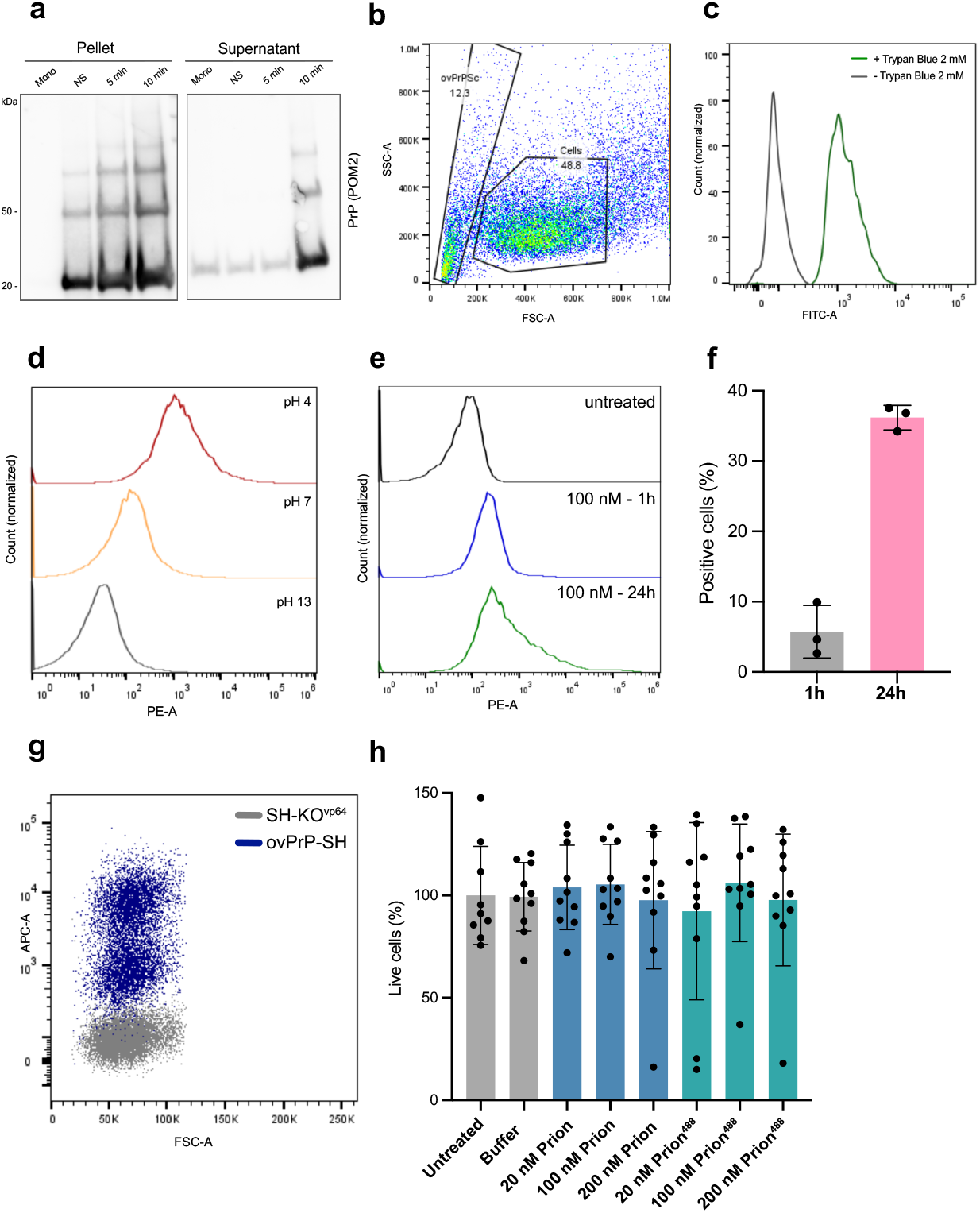
**a.** Western Blots of pellet and supernatant fractions from a density cushion with 60% iodixanol. Insoluble prion^488^ aggregates were present mostly in the pellet fraction. After 10 minutes of sonication, a signal was visible also in the supernatant fraction. Mono: monomeric ovine PrP; NS: prion^488^ aggregates not sonicated; 5 min and 10 min: prion^488^ aggregates sonicated for 5 min and 10 min, respectively. **b.** Representative flow cytometry dot plots showing forward scatter area (FSC-A) versus side scatter area (SSC-A) profiles of prion^488^ aggregates and untreated cells. The two samples were mixed immediately before acquisition. **c.** Histogram showing fluorescence intensity of prion^488^ aggregates (X axis, FITC channel) acquired without and with the addition of 2 mM Trypan Blue. Fluorescence intensity decreases almost to zero when Trypan Blue is present in the sample during acquisition. **d.** Histograms of fluorescence intensity of Prion^pHrodo^ acquired in PBS at pH 4, pH 7 and pH 13. Fluorescence intensity of prion^pHrodo^ decreases at higher pH. **e.** Histograms of fluorescent intensity of cells exposed to 100 nM prion^pHrodo^ (1h and 24 h) compared to untreated cells. **f.** Uptake quantification of prion^pHrodo^ after 1h and 24h incubation. Dots represent single wells treated individually with prion^pHrodo^ preparation. **g.** Surface staining of ovPrP on SH-KO^vp64^ cells and ovPrP-SH cells transduced with a construct for ovPrP expression. Staining was performed on live cells using AF647-conjugated 6D11 antibody. **h.** Viability assay of SH-KO^vp64^ upon incubation with increasing concentrations of unlabelled prion aggregates and prion^488^ aggregates for 24 hours. Results are expressed as percentage of live cells normalized on the mean of the untreated controls. Dots represent single wells treated individually with prion^488^ preparations in n=2 independent epxeriments. Statistical test: Brown-Forsythe ANOVA with Dunnett’s T3 post-hoc test. Error bars: mean ± SD.

**Supplementary Figure 3.**
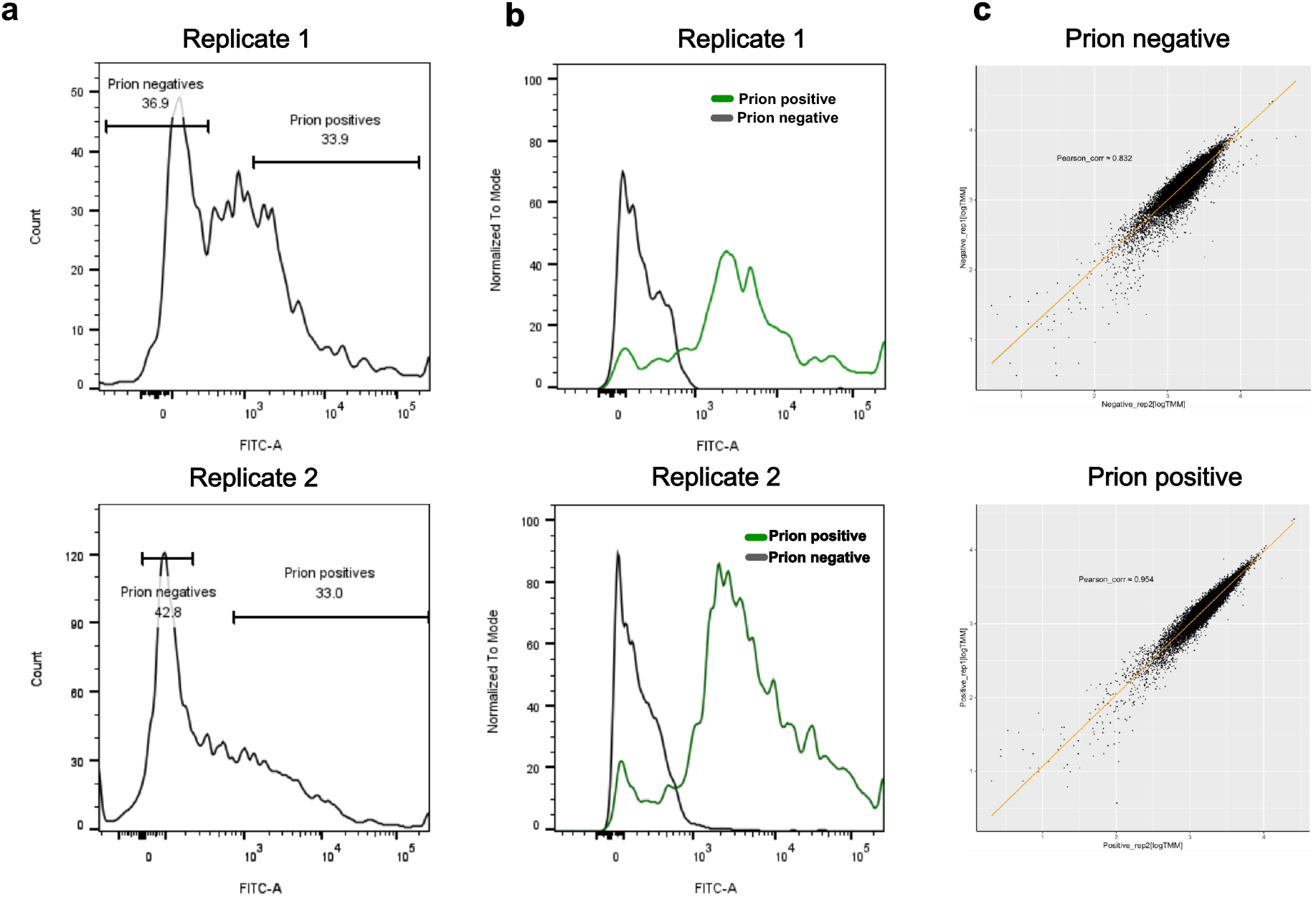
**a.** Histogram plots showing the FITC fluorescence intensity profile of the cellular population treated with Prion^488^ aggregates for 24 hours, immediately before sorting. The two plots refer to the first (left) and second (right) biological replicate of the primary screen. Horizontal lines represent the two sorted populations. **b.** Reanalysis of the two sorted populations in the first (left) and second (right) biological replicate of the primary screen. **c.** Replicate correlation for the prion negative (left, Pearson corr.:0.83)) and prion positive (right, Pearson corr.:0.95) populations, calculated on NGS read counts.

**Supplementary Figure 4.**
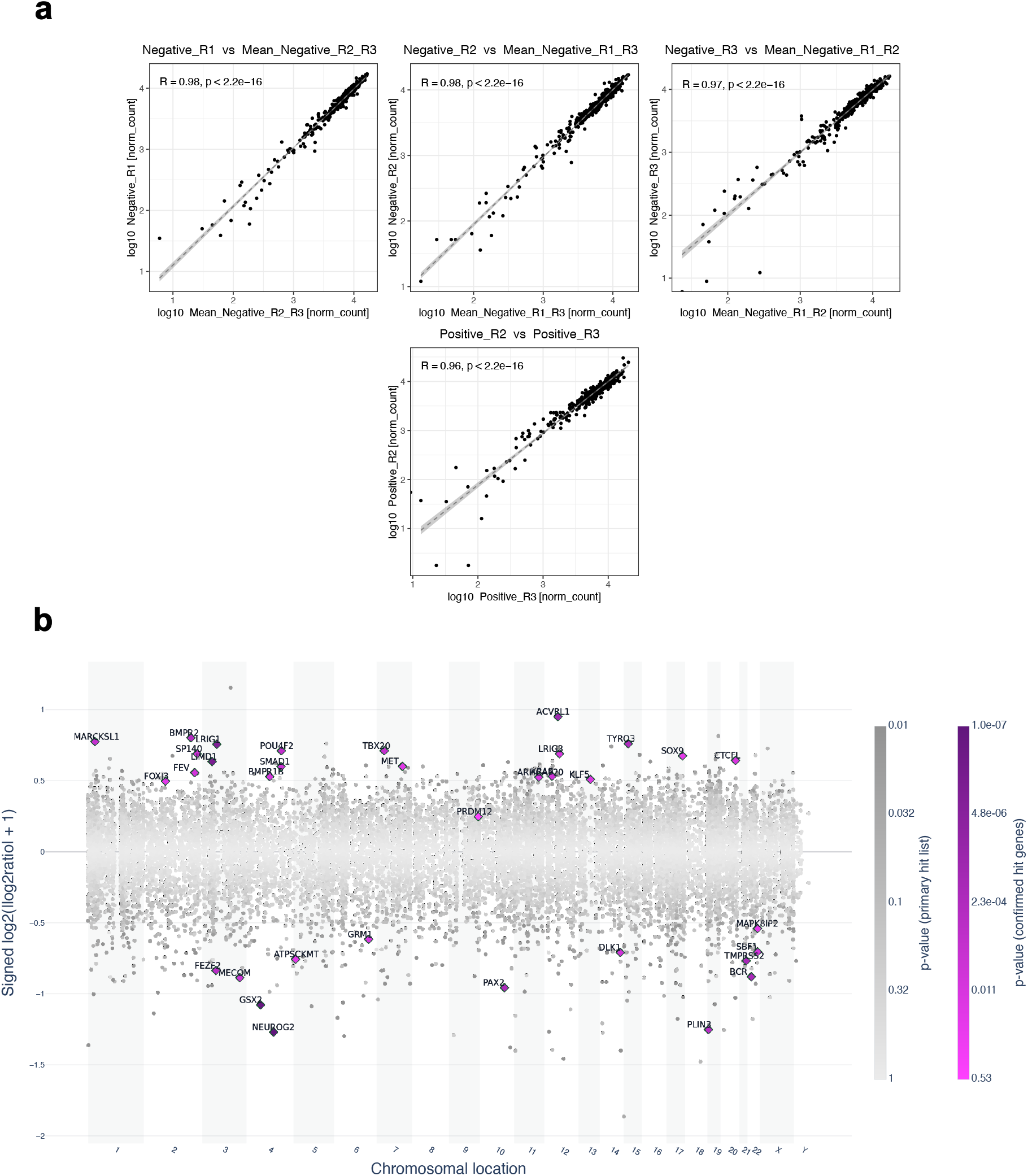
**a.** Correlation analysis of biological replicates for the prion-negative (top) and prion^+^ (bottom) populations in the secondary screen. Correlations were calculated based on NGS read counts. Each of the three biological replicates was compared with the other two. The prion^488+^ population from the first replicate showed poor correlation with the other two and was therefore excluded from further analysis. **b.** Plot showing log_2_FC (y-axis) and p-value (color code) of confirmed hit genes plotted against their genomic coordinates (x-axis).

**Supplementary Figure 5.**
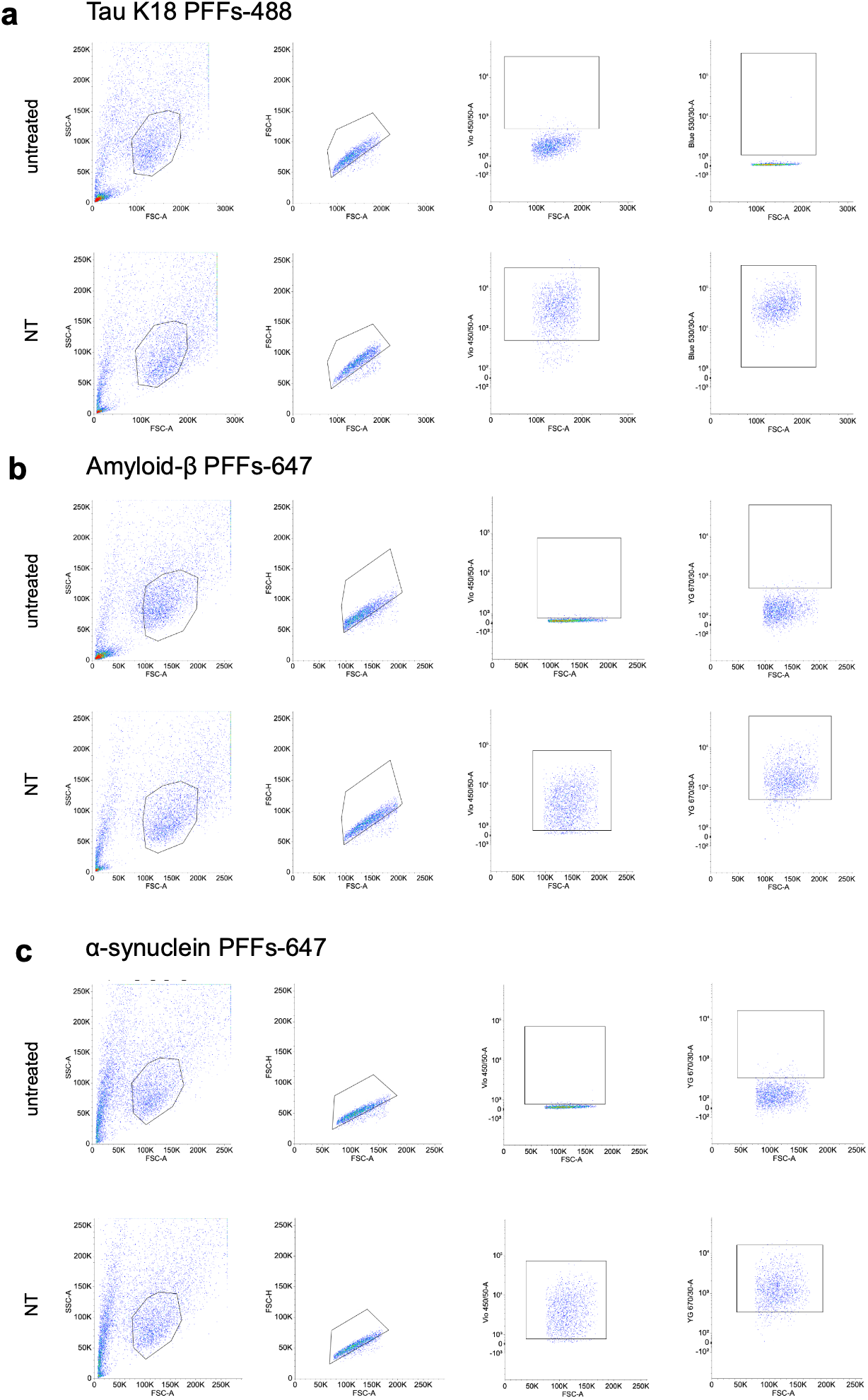
Gating strategy for flow cytometry-based uptake assays of Tau K18-488 (a), Amyloid β-647 (b) and α synuclein-647 (c) pre-formed fibrils (PFFs). Cells were gated based on FSC-A and SSC-A, and single cells were gated based on FSC-A and FSC-H values. qgRNA-transduced cells were gated based on the expression of BFP protein. Probe-positive cells were gated based on untreated controls.

**Supplementary Figure 6.**
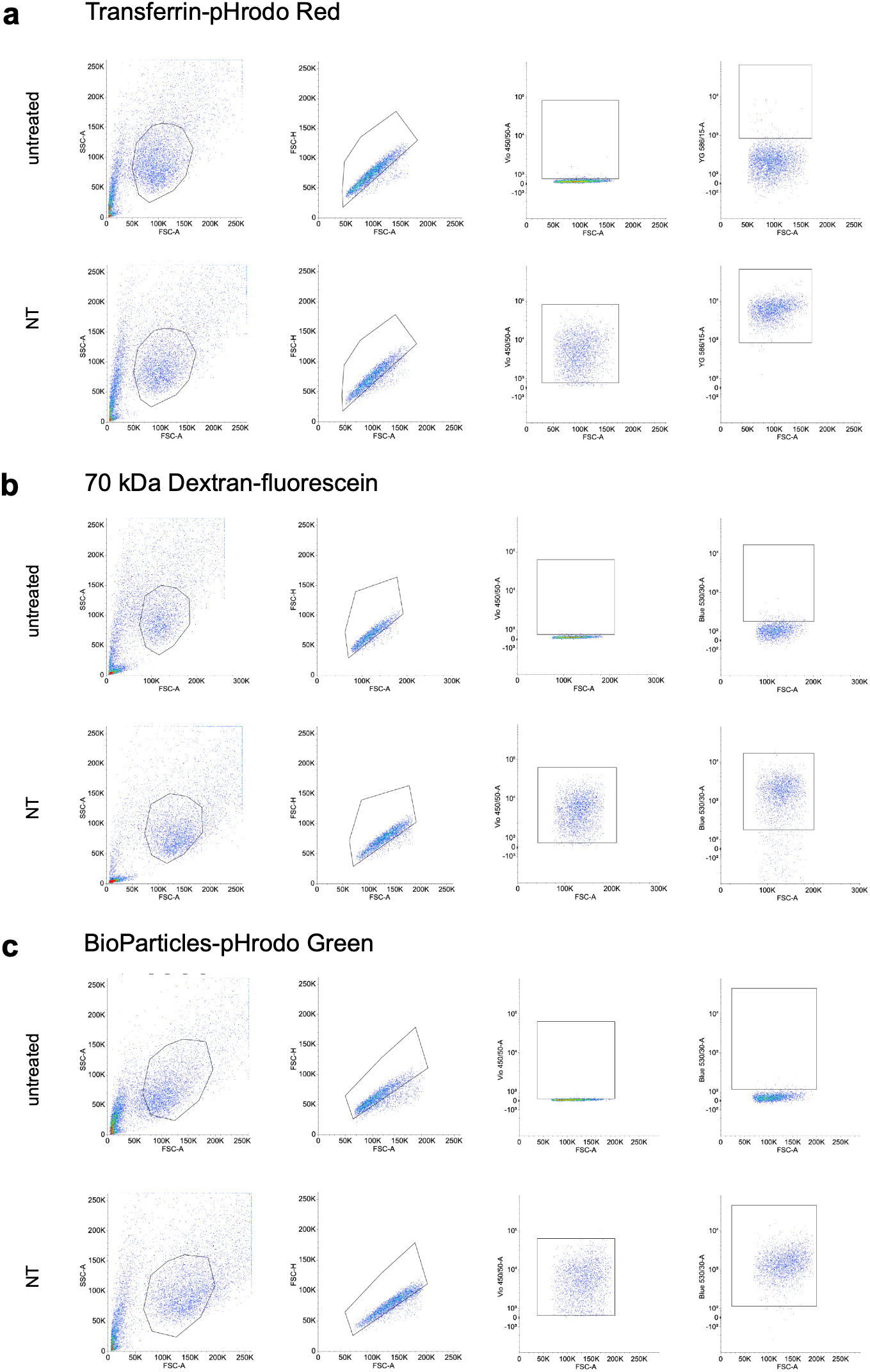
Gating strategy for flow cytometry-based uptake assays of transferrin-pHrodo Red (a), 70 kDa dextran-fluorescein (b) and BioParticles-pHrodo Green (c). Cells were gated based on FSC-A and SSC-A, and single cells were gated based on FSC-A and FSC-H values. qgRNA-transduced cells were gated based on the expression of BFP protein. Probe-positive cells were gated based on untreated controls.

**Supplementary Figure 7.**
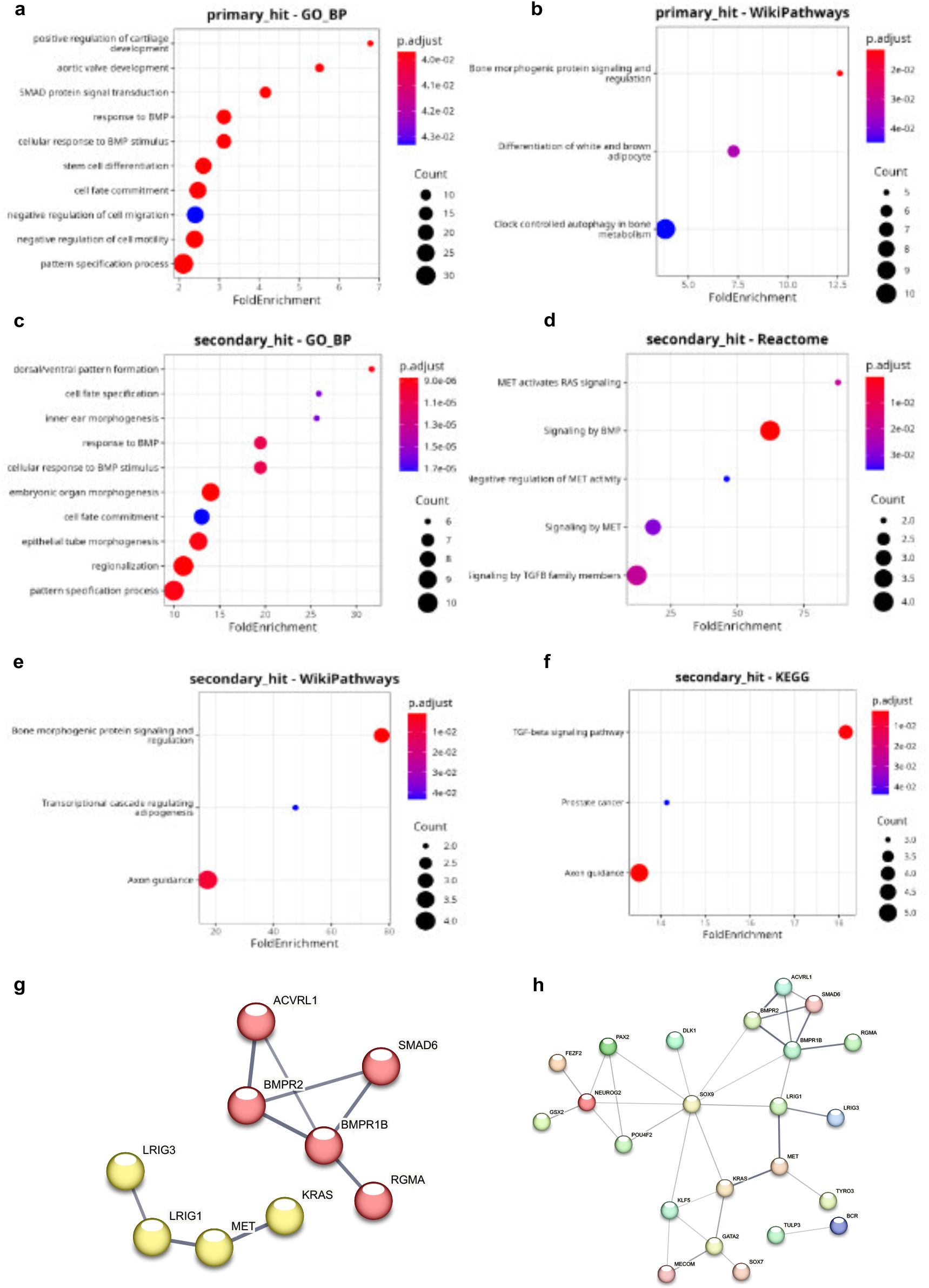
a-b. Over Representation Analysis (ORA) of primary screen hitlist using the GO:BP (a) and WikiPathways (b) databases. **c-f.** ORA of confirmed hit list from the secondary screen using the GO:BP (c), Reactome (d), WikiPathways (e) and KEGG (f) databases. All ORA are visualized using Cluster Profiler. **g-h**. PPI network of confirmed hit genes. The network was generated using STRING online tool. Nodes represent proteins, and edges represent known or predicted. Confidence score cutoff ≥ 0.7 (high, g) and ≥ 0.4 (moderate, h).

**Supplementary Figure 8.**
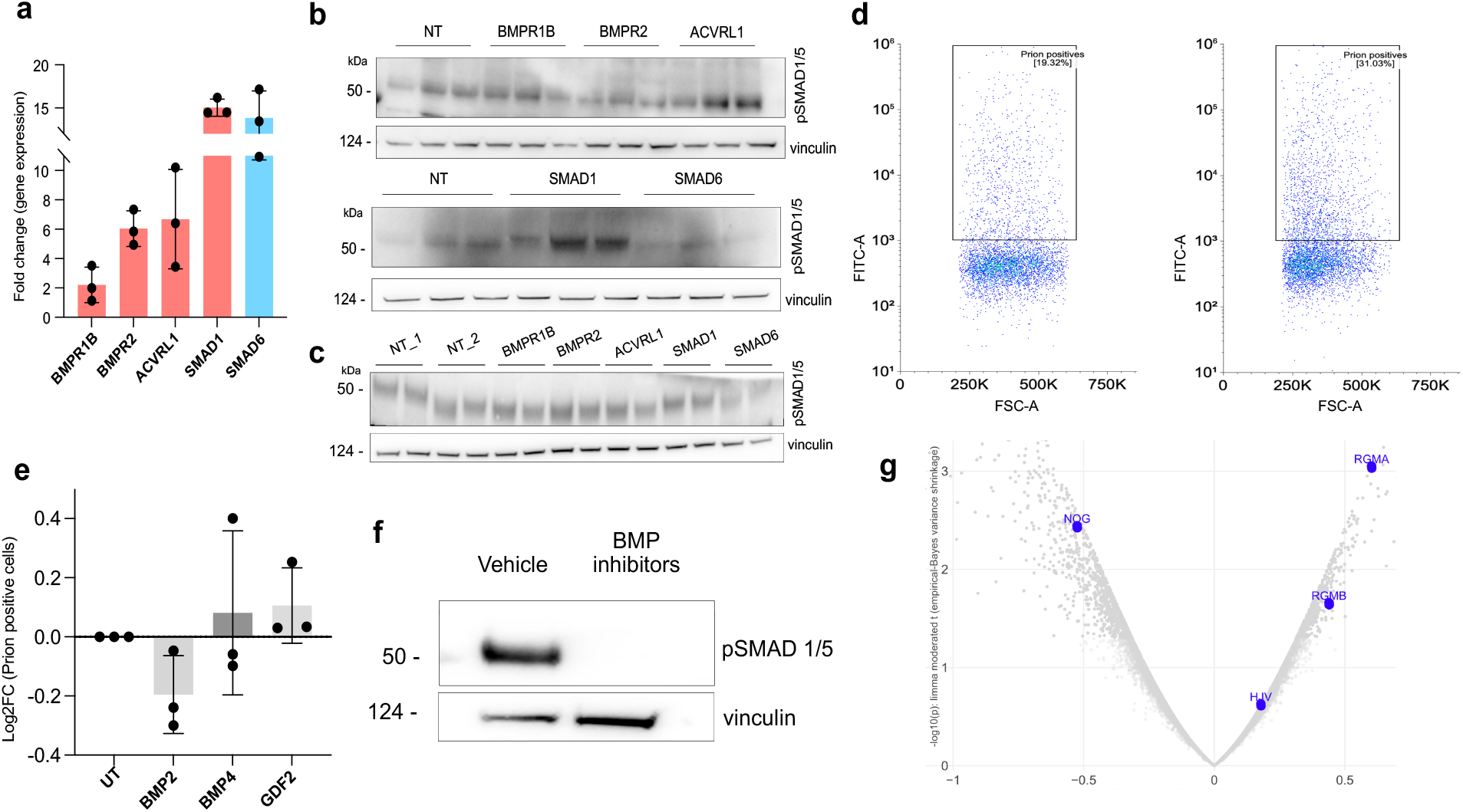
**a.** qPCR analysis of the expression levels of BMP hit genes upon genetic upregulation using qgRNA guides. Activation was measured 6 days after infection. Data are shown as fold change calculated on a non-targeting control condition. Each dot is the average of three technical replicates in n=3 independent experiments. **b-c.** Western Blots for pSMAD1/5 upon genetic upregulation of BMP hit genes. Vinculin was used as loading control. Quantification is shown in Fig. 4d-e. **D.** Representative flow cytometry dot plots of Prion^488^ uptake in SH-KO^vp64^ cells treated with a combination of BMP peptides. Quantification is shown in Fig. 4g. **e.** Quantification of prion^488^ uptake in SH-KO^vp64^ cells treated with individual BMP peptides. Dots represent individual (n = 3) experiments and are the average of three technical replicates. **f.** Western Blot showing pSMAD1/5 levels in SH-KO^vp64^ treated with BMP inhibitors or with vehicle as control. Vinculin: loading control. **g.** Volcano plot displaying log_2_FC and p-values of all protein-coding genes in the primary screen; *NOG* (encoding for noggin), RGMA, RGMB and HJV are highlighted. Error bars: mean ± SD.

**Supplementary Figure 9.**
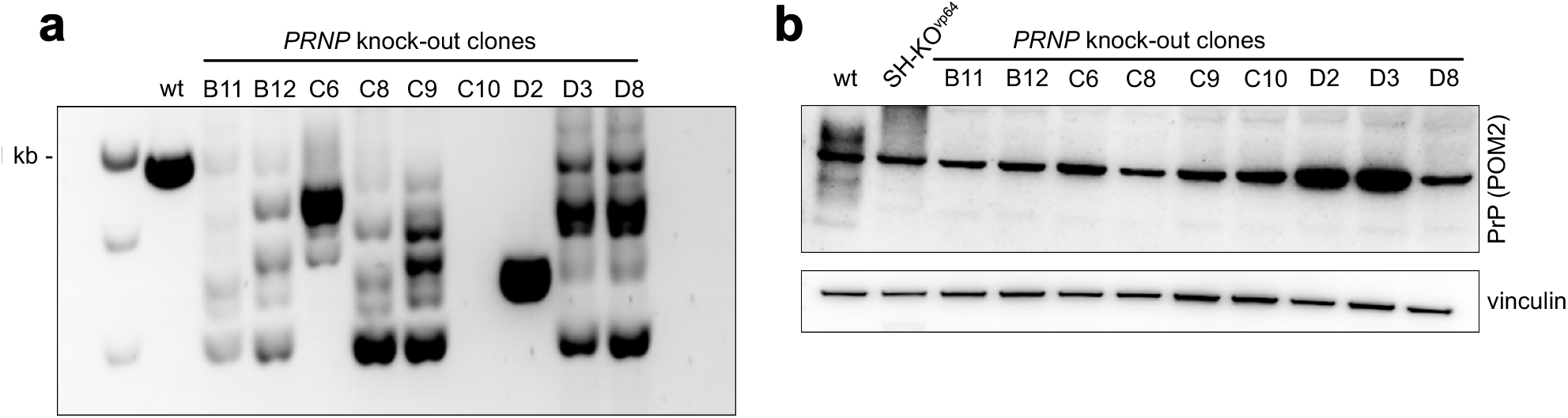
**a.** Agarose gel showing genotyping PCR of single HEK293A clones treated with qgRNA targeting human PRNP to confirm PRNP knockout. Wild-type HEK293A were used as positive control. **b.** Western blot showing PrPC levels in different HEK293A monoclones treated with qgRNA targeting human PRNP. Wild-type HEK293A and SHSY-5Y cells with knockout of PRNP are used as positive and negative controls, respectively. Vinculin was used as loading control.

**Supplementary Figure 10.**
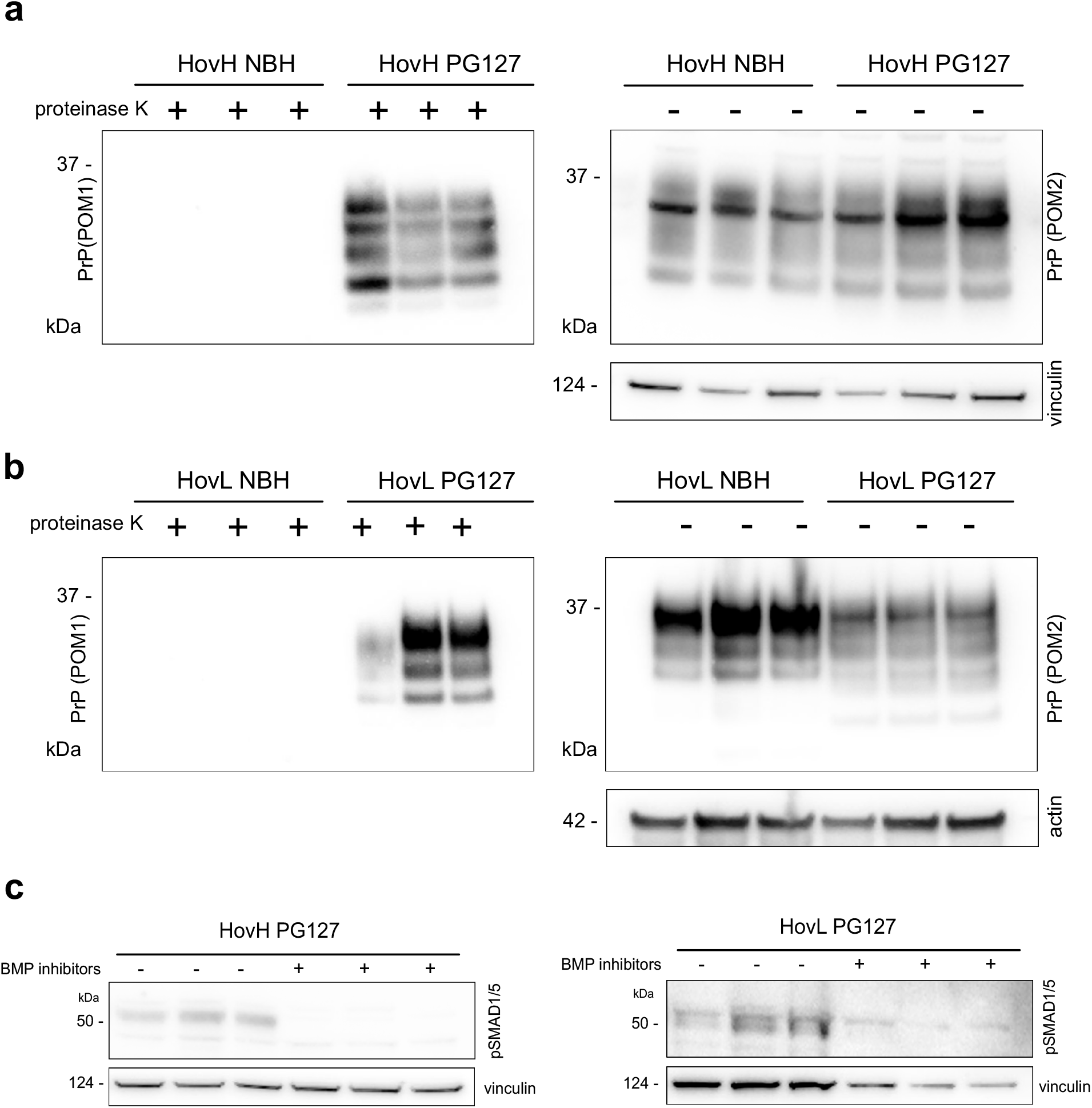
a-b. Western Blots of HovH (**a**) and HovL (**b**) cell lines infected either with non-infected (NBH) or prion-infected (PG127) brain homogenates. Proteinase K digestion shows the presence of PK-resistant PrP^Sc^ only in the PG-infected cells. Vinculin (**a**) and actin (**b**) were used as loading controls. **c.** Western Blots showing pSMAD1/5 levels in HovH and HovL treated with BMP inhibitors and vehicle as control. Vinculin was used as loading control.

