## Supplemental Document 3 for "Control of Prion Uptake by Bone Morphogenetic Protein Signaling" (from another version of this preprint: version 2, 2026-04-11)

#### Supplement: Derivation of the 209-gene human BMP signaling gene catalog and comparison to published genetic data on neurodegenerative diseases

The curated backbone came from the human Reactome BMP pathway and the current human MSigDB collections, which expose both curated Reactome pathway gene sets and GO-derived ontology gene sets. Reactome describes canonical BMP signaling through type I and type II serine/threonine kinase receptors, SMAD1/5/8, and SMAD4, with modulators such as endoglin and RGM-family factors [1,4]. The MSigDB overview page identifies the current human release as v2026.1.Hs, and the relevant ontology gene sets specify their own GO release history on the individual gene-set pages [2-9].

##### 2.2 Formal inclusion rule

| Evidence bucket | External source | Version / exact source | Source size | Role in released derivation |
|---|---|---|---|---|
| Reactome_core | REACTOME_SIGNALING_BY_BMP / Reactome Signaling by BMP | MSigDB 2026.1.Hs; gene set updated to Reactome 95 [2,4] | 31 | Inclusion source |
| GO_response | GOBP_RESPONSE_TO_BMP | GO:0071772; MSigDB 2025.1.Hs, GO Release 2025-03-16 [5] | 193 | Inclusion source |
| GO_reg | GOBP_REGULATION_OF_BMP_SIGNALING_PATHWAY | GO:0030510; MSigDB 2025.1.Hs, GO Release 2025-03-16 [6] | 127 | Inclusion source |
| GO_pos_reg | GOBP_POSITIVE_REGULATION_OF_BMP_SIGNALING_PATHWAY | GO:0030513; MSigDB 2025.1.Hs, GO Release 2025-03-16 [7] | 40 | Inclusion source |
| GO_receptor_act | GOMF_BMP_RECEPTOR_ACTIVITY | GO:0098821; MSigDB 2026.1.Hs, GO Release 2025-10-10 [9] | 8 | Inclusion source |
| GO_receptor_bind | GOMF_BMP_RECEPTOR_BINDING | GO:0070700; human MSigDB download endpoint accessed 2026-03-30; GO term definition in AmiGO [10] | 13 | Inclusion source |
| Lit_direct | Manual literature additions | Eight genes added explicitly from primary/review literature [11-16] | 8 | Inclusion source |
| GO_neg_reg | GOBP_NEGATIVE_REGULATION_OF_BMP_SIGNALING_PATHWAY | GO:0030514; MSigDB 2025.1.Hs, GO Release 2025-03-16 [8] | 75 | Corroborative overlay only; not required to reproduce the 209-gene inclusion universe |

The catalog was constructed by combining, as they existed at the time of access, all human gene symbols from the Reactome core BMP pathway, the GO/MSigDB gene sets for BMP response, BMP regulation, positive regulation of BMP signaling, BMP receptor activity, and BMP receptor binding, together with the predefined literature-supported additions DEK, EIF4A2, HAMP, HEY1, ID2, ID3, JUNB, and SP7, and then removing duplicates to yield 209 unique genes.

Deduplication was performed at the released gene-symbol level, with one retained row per symbol. No broad synonym expansion, family-level roll-up, or cross-species harmonization was applied beyond using the human symbols exposed by the selected resources and noting a few aliases in the notes field.

The negative-regulation source *GOBP_NEGATIVE_REGULATION_OF_BMP_SIGNALING_PATHWAY* [8] was BMP-relevant and therefore reviewed, but it is not required to reproduce the 209-row inclusion universe. In the released artifact it functions only as a corroborative bucket on five already retained genes (BMPER, ERFE, GPR155, HJV, and TMPRSS6).

The source mix was deliberately broad enough to capture pathway-core genes, extracellular modulators, intracellular regulators, direct targets, and downstream/contextual responders. Reactome contributed canonical pathway structure [1,4]; GO response/regulation terms broadened the universe to include downstream and modulatory genes [5-8]; receptor-activity and receptor-binding terms captured mechanistic receptor-side or ligand-side relations [9,10]; and the literature-only set added well-known direct or early targets that were not reliably present in the selected ontology sets [11-16].

#### 3. Column-by-column legend for the spreadsheet

| Column | Meaning in the released table |
|---|---|
| gene_symbol | Row identifier. Deduplication was performed at the released gene-symbol level, with one row per retained symbol. No broad synonym collapse was applied beyond noting a few common aliases in the notes field. |
| Roles | Manual functional classification of the retained gene in the context of BMP signaling. Multiple roles are separated by a semicolon and indicate that the release preserved dual interpretation rather than forcing a single class. |
| confidence_index_1to5 | Ordinal confidence score. In this release, only values 2, 3, 4, and 5 are used. The score is a curated robustness estimate, not a probability, p-value, citation count, or effect-size measure. |
| confidence_label | Verbal form of the ordinal score: 5 = very high; 4 = high; 3 = moderate; 2 = low-to-moderate. |
| evidence_buckets | Semicolon-delimited summary of source categories supporting the row. This field is a presentation summary and not a complete incidence matrix of every overlapping source membership. |
| evidence_bucket_count | Count of tokens present in evidence_buckets. It counts displayed buckets only; it does not count studies, experiments, or all possible overlapping database memberships. |
| brief_basis | Exact one-sentence rationale archetype used to drive the confidence score. In the released artifact, the confidence score can be reproduced exactly from this field. |
| representative_source_urls | Representative, not exhaustive, supporting URLs. The list is illustrative and sufficient for traceability, but it is not intended to be a full bibliography for each gene. |

#### 4. Functional role assignment

The role labels were chosen to map the heterogeneous source evidence into a stable, publication-friendly vocabulary. When the evidence supported more than one role, we preserved both by separating them with a semicolon.

| Role label | Operational definition used in the release |
|---|---|
| ligand | Secreted BMP-family ligand or closely BMP-related TGFB-family ligand treated as part of the BMP signaling universe in curated pathway/ontology resources. |
| receptor | Canonical type I or type II BMP receptor, or receptor explicitly annotated as having BMP receptor activity in curated resources. |
| co-receptor / receptor-associated factor | Surface or receptor-proximal factor that changes ligand-receptor engagement or signaling competence but is not itself the canonical signaling receptor. |

| Role label | Operational definition used in the release |
|---|---|
| extracellular modulator | Extracellular antagonist, agonist, scaffold, or binding factor that changes BMP ligand availability, sequestration, presentation, or receptor access. |
| intracellular mediator/regulator | Intracellular transducer or regulator of pathway transmission, attenuation, trafficking, ubiquitylation, phosphatase control, or transcriptional competence. |
| direct target gene | Gene treated in the release as a direct or early primary BMP-responsive transcriptional target, based on literature or repeated review-level recognition. |
| secondary/context-dependent response gene | Gene retained because it recurs as a downstream transcriptional response, differentiation marker, or context-specific BMP-responsive gene, without being treated as a universal pathway-core component. |
| ncRNA regulator | microRNA, lncRNA, or antisense RNA retained because it appears in BMP response/regulation sources or because the release treated it as BMP-regulatory in a recurrent manner. |
| context-dependent BMP-associated gene | Residual category used where a BMP link was retained but a more specific mechanistic class would overstate the evidence. In this release, this category is used only once (PYCARD). |

#### 5. Computation of confidence scores

Confidence was assigned by a **rule-based ordinal rubric**. The release first assigned each gene to one of a small number of textual rationale archetypes recorded verbatim in **brief_basis**. Each archetype was then mapped to a fixed integer score. Because the mapping is one-to-one, the score can be reproduced exactly from the brief_basis field.

The rubric privileged **mechanistic proximity** and **evidence provenance**. Canonical receptors, canonical ligands, core SMAD mediators, and repeatedly validated direct targets received the highest scores. Strong modulators, co-receptors, intracellular regulators, and context-specific but well-supported direct targets generally received score 4. Recurrent but context-dependent curated response/regulation genes received score 3. Narrower or less BMP-specific ontology-level associations received score 2.

**Evidence_bucket_count** is descriptive only. It helped summarize provenance, but it was not the scoring formula. A gene with one highly specific source (for example a canonical receptor or a literature-validated direct target) could score higher than a gene appearing in multiple broad ontology sets.

##### 5.2 Exact mapping used in the released artifact

| Exact brief_basis text in released sheet | Score | Interpretation |
|---|---|---|
| Canonical BMP ligand/receptor connection in curated pathway resources. | 5 (very high) | Canonical/core pathway membership or repeatedly validated direct BMP target. |
| Core BMP signal-transduction component with strong curated support. | 5 (very high) | Canonical/core pathway membership or repeatedly validated direct BMP target. |
| Repeatedly validated direct BMP target in review/primary literature. | 5 (very high) | Canonical/core pathway membership or repeatedly validated direct BMP target. |
| BMP-family or BMP-related ligand with strong but somewhat context-specific pathway support. | 4 (high) | Strong BMP connection, but not as universally canonical as the score-5 core. |
| Strong literature support as a direct BMP-responsive target in defined contexts. | 4 (high) | Strong BMP connection, but not as universally canonical as the score-5 core. |
| Well-supported BMP modulator/co-receptor in curated pathway or GO resources. | 4 (high) | Strong BMP connection, but not as universally canonical as the score-5 core. |
| Well-supported intracellular BMP regulator in curated resources. | 4 (high) | Strong BMP connection, but not as universally canonical as the score-5 core. |
| Primary-literature BMP response gene in specific differentiation contexts; likely downstream rather than pathway-core. | 3 (moderate) | Recurrent and defensible BMP association, but materially context dependent. |
| Recurrent BMP response/regulation annotation across curated resources; context dependence remains important. | 3 (moderate) | Recurrent and defensible BMP association, but materially context dependent. |

| Exact brief_basis text in released sheet | Score | Interpretation |
|---|---|---|
| Recurrent ncRNA-level BMP response/regulation annotation; mechanism remains context dependent. | 3 (moderate) | Recurrent and defensible BMP association, but materially context dependent. |
| Limited or mainly context-specific BMP evidence relative to the canonical pathway core. | 2 (low-to-moderate) | Retained BMP association with lower specificity or narrower context. |
| Reported mainly in GO response/regulation annotations; direct BMP specificity is less robust. | 2 (low-to-moderate) | Retained BMP association with lower specificity or narrower context. |

##### 5.3 Distribution in the current release

| Metric | Count |
|---|---|
| Unique released gene symbols | 209 |
| Rows with confidence 5 (very high) | 27 |
| Rows with confidence 4 (high) | 68 |
| Rows with confidence 3 (moderate) | 70 |
| Rows with confidence 2 (low-to-moderate) | 44 |
| Rows with confidence >= 4 ("High_confidence_core" sheet) | 95 |

No score 1 rows are present in the current version. A score of 1 was effectively reserved for links too weak or too anecdotal to retain, and therefore no such rows survived into the released catalog.

#### 6. Audit findings and transparency notes

The central audit result is positive: the released 209-gene inclusion universe is exactly reproducible from the set union defined in Section 2. However, one field in the spreadsheet requires a publication note for precision. The **evidence_buckets** column behaves as a concise display field rather than as a lossless provenance matrix. Two small compressions were observed during audit:

| Audit item | Rows affected | Effect on inclusion universe |
|---|---|---|
| GO_response membership present in source but not explicitly listed in evidence_buckets (HIPK2) | 1 | None; row retained and counted in the 209-gene union |
| GO_reg membership present in source but not explicitly listed in evidence_buckets (BMP4, SFRP1, SFRP2, SKOR2, SKP1, SMURF1, SORL1, SOX11, TBX20, UBE2O) | 10 | None; rows retained and counted in the 209-gene union |
| GO_neg_reg used only as corroborative overlay (BMPER, ERFE, GPR155, HJV, TMPRSS6) | 5 | None; this set is not needed to reproduce the final 209 rows |

**These compressions do *not*** alter row retention, role assignment, or confidence scoring. They matter only for anyone attempting to interpret evidence_buckets or evidence_bucket_count as a complete source-membership matrix. For publication, the safest wording is that the field is a **summary provenance field**, not an exhaustive encoding of all overlaps.

#### 7. Appendix A - literature-only additions that expanded the inclusion universe

The eight genes below are the only additions required beyond the six inclusion-source buckets to reproduce the released 209 rows. Their bucket label **Lit_direct** should be read as a historical shorthand for "manually added from the literature" rather than as a guarantee that every such gene is a universally direct target in every context.

| Gene | Released role | Reason this gene was explicitly retained | Representative literature |
|---|---|---|---|
| DEK | direct target gene | Added as a literature-only direct BMP target not captured by the selected ontology/pathway union. | Hollnagel et al., 1999 [11] |
| EIF4A2 | direct target gene | Represents the eIF4AII literature target reported in embryonic stem-cell BMP induction experiments. | Hollnagel et al., 1999 [11] |

| Gene | Released role | Reason this gene was explicitly retained | Representative literature |
|---|---|---|---|
| HAMP | direct target gene | Included as a strong BMP/HJV/SMAD-responsive effector with promoter-level BMP responsiveness and extensive liver BMP-SMAD support. | Truksa et al., 2009 [15]; Silvestri et al., 2019 [16] |
| HEY1 | direct target gene | Included as an early/direct BMP9-SMAD target with promoter/chromatin evidence in osteogenic contexts. | Sharff et al., 2009 [14] |
| ID2 | direct target gene | Literature-only direct BMP target retained despite absence from the selected ontology inclusion sets. | Hollnagel et al., 1999 [11]; Borok et al., 2020 [12] |
| ID3 | direct target gene | Literature-only direct BMP target retained despite absence from the selected ontology inclusion sets. | Hollnagel et al., 1999 [11]; Borok et al., 2020 [12] |
| JUNB | secondary/context-dependent response gene | Rapid early BMP response retained from primary literature, but not elevated to universal direct-target status in the release. | Hollnagel et al., 1999 [11] |
| SP7 | secondary/context-dependent response gene | BMP-induced osteogenic downstream effector retained from primary literature; not treated as direct because induction requires new protein synthesis. | Lee et al., 2003 [13] |

#### 8. Appendix B - role distribution in the released table

| Role label | Count |
|---|---|
| secondary/context-dependent response gene | 59 |
| intracellular mediator/regulator | 45 |
| extracellular modulator | 35 |
| ligand | 18 |
| ncRNA regulator | 18 |
| direct target gene | 12 |
| co-receptor / receptor-associated factor; extracellular modulator | 11 |
| receptor | 8 |
| intracellular mediator/regulator; direct target gene | 2 |
| context-dependent BMP-associated gene | 1 |

### Supplementary Methods: Interrogation of AD, ALS and PD human-genetics datasets for enrichment of BMP-related genes

#### Datasets interrogated

The disease comparisons were intentionally anchored to the largest or most informative human-genetics studies discussed in this project, with the primary analysis layer chosen to match the question of pathway overrepresentation. Common-variant GWAS/MAGMA studies were used as the primary evidence for AD and PD, whereas the largest harmonized rare-variant exome study was used as the primary evidence for ALS.

| Disease | Primary study used | Study size / modality | Outputs interrogated for BMP audit |
|---|---|---|---|
| **AD** | Bellenguez et al., 2022 [1] | 111,326 clinically diagnosed/proxy cases; 677,663 controls; common-variant GWAS with MAGMA pathway analysis | Corrected MAGMA gene-set results (10,271 sets), pathway/cell-type interpretation, and the prioritized genes reported for the newly identified loci. |
| **ALS** | Hop et al., 2026 [2] | 17,919 ALS cases; 200,703 controls across discovery and replication; rare-variant exome single-variant and URV burden analyses | Geneset burden results (11,777 GO/KEGG/Reactome sets), exome-wide significant genes/variants, replicated candidate genes, and candidate-gene GO follow-up. |
| **PD** | Kim et al., 2024 [3] | 49,049 PD cases; approximately 18.6k proxy cases; 2,458,063 controls; multi-ancestry GWAS with MAGMA/FUMA analysis | Corrected MAGMA/FUMA ontology results (16,992 sets), 12 novel loci, 6 single-variant fine-mapped loci, and 25 putative genes nominated from the novel loci. |
| **PD (secondary orthogonal checks)** | Hop et al., 2024 [4]; Pitz et al., 2024 [5] | Familial exome rare-variant study and large population-level rare-variant meta-analysis | Highlighted significant genes/variants only; these studies were used as confirmatory gene-level checks, not as the primary pathway-level test. |

##### Exact public outputs queried

- **AD / Bellenguez 2022.** Main article pathway-analysis section; Supplementary Table 15 (pathway enrichment); Supplementary Tables 16–18 (cell-type and pathway-interaction analyses); Fig. 2a and Supplementary Tables 19–30 for gene prioritization; GWAS Catalog accession GCST90027158.

- **ALS / Hop 2026.** Main article results section on geneset burden; Table 1 and corresponding figures for single-variant hits; Supplementary Data 1 (single-variant associations), Supplementary Data 3 (ultrarare gene-based analyses), Supplementary Data 7–8 (geneset burden), and Supplementary Data 11 (full list of GO terms for candidate genes).

- **PD / Kim 2024.** Main article gene-set analysis section; Supplementary Table 9 (significant and conditional ontology terms); Table 3 and locus-level results for fine-mapping; the reported 12 novel loci, 6 single-variant fine-mapped loci, and 25 putative genes nominated by eQTL/SMR; GWAS Catalog accession GCST90275127 and the GP2 analysis-code archive.

- **PD secondary checks.** Hop 2024 familial PD exome letter and Pitz 2024 rare-variant meta-analysis; only the studies' highlighted significant genes and variants were used, because these papers were employed as orthogonal gene-level checks rather than as the primary pathway-level evidence.

Note on the PD paper: the Kim et al. article contains a minor internal discrepancy in the number of proxy cases (18,785 in the abstract versus 18,618 in the main text). This did not affect the comparison procedure, because the PD study was treated as a single predefined dataset and the audit relied on the study authors' own pathway and locus outputs rather than on recalculation from raw case counts.

#### Common comparison framework

**The comparison was performed within each disease study's own inferential framework.** This was essential because the three diseases were represented by different classes of genetic study: common-variant GWAS with MAGMA gene-set testing for AD and PD, and rare-variant exome single-variant plus ultrarare-variant (URV) burden analyses for ALS. No attempt was made to pool raw P values across diseases.

- **Fixed BMP input.** The 209-gene human BMP catalog was frozen before any disease-specific review. No disease-specific additions, deletions or family-level roll-up were allowed during the comparison.

- **Primary evidence layer: study-internal pathway significance.** A disease dataset was considered supportive of BMP overrepresentation only if the study's own corrected pathway or geneset analysis identified at least one explicitly BMP-related term. For AD and PD, this meant the authors' MAGMA competitive gene-set results after q-value/FDR correction. For ALS, this meant the authors' rare-variant geneset burden results after the study-specific filtering used to remove single-gene-driven terms.

- **Secondary evidence layer: symbol-level gene overlap.** The authors' reported disease genes—such as prioritized genes, lead-locus genes, fine-mapped genes, replicated rare-variant genes, or top genes inside significant disease pathways—were intersected with the fixed 209 BMP genes. This secondary layer was used as a confirmatory audit only; isolated gene overlaps were not allowed to override a negative pathway-level result.

- **BMP specificity rule.** Generic TGF-β, cytokine or developmental terms were not counted as BMP-positive unless the pathway label or the authors' interpretation was explicitly BMP-specific. In practice, a pathway was counted as BMP-related only if it contained the phrase "BMP", "bone morphogenetic", "signaling by BMP", "BMP receptor", "response to BMP", or "regulation of BMP signaling".

- **Symbol harmonization.** Gene symbols were uppercased and matched against approved human gene symbols. Where a source used an obvious historical alias or protein synonym, it was mapped back to the approved symbol before overlap was scored. Duplicates were removed after normalization.

#### Exact step-by-step workflow

1. Lock the 209-gene BMP catalog defined in the companion document and use it unchanged for all disease comparisons.

2. Select the prespecified disease dataset to be interrogated (Bellenguez 2022 for AD; Hop 2026 for ALS; Kim 2024 for PD; Hop 2024 and Pitz 2024 as secondary PD rare-variant checks).

3. Retrieve the study outputs that the authors themselves used to summarize disease signal: corrected pathway/gene-set analyses, prioritized genes, lead-locus genes, fine-mapped genes, burden hits, replicated candidate genes, and any disease-relevant GO or pathway follow-up tables explicitly reported by the authors.

4. Perform a pathway-name scan of the corrected pathway results using the prespecified BMP keyword set: BMP; bone morphogenetic; signaling by BMP; BMP signaling; BMP receptor; response to BMP; regulation of BMP signaling.

5. Perform a symbol-level overlap audit by exact matching the fixed 209 BMP genes to the study's reported disease-gene lists after symbol normalization.

6. Record pathway-level and gene-level findings separately. Pathway-level negative means that no study-corrected pathway was explicitly BMP-related. Gene-level negative means that the overlap audit did not reveal a concentration of BMP genes among the disease-associated or prioritized genes highlighted by the paper.

7. Call the disease dataset "not supportive of BMP overrepresentation" only when both layers were negative: no corrected BMP pathway and no convincing gene-level concentration of BMP genes.

#### Disease-specific implementation

##### Alzheimer's disease (AD)

The AD analysis used Bellenguez et al. as the primary dataset because it is a large two-stage AD/ADRD GWAS with explicit pathway analysis and public summary-statistics deposition. The study reported 75 loci and performed MAGMA competitive gene-set analysis over 10,271 gene sets, using q values to control for multiple testing.

The BMP audit was done in two layers. First, the corrected MAGMA pathway outputs and the accompanying pathway interpretation were screened for any explicitly BMP-related term. Because the Bellenguez paper already used the competitive MAGMA framework recommended for pathway enrichment, this published, study-internal result was treated as the decisive enrichment layer for AD. Second, the prioritized disease genes reported by the authors—especially the 55 prioritized genes from the newly identified loci—were intersected with the fixed BMP catalog after symbol normalization.

AD was classified as not supportive of BMP overrepresentation when no significant pathway term was BMP-specific and the gene-prioritization layer likewise failed to reveal a coherent BMP cluster. Non-BMP signals highlighted by the authors, such as amyloid/tau biology, microglial expression, innate immunity, endocytosis and TNF/LUBAC-related interpretation, were recorded as disease-relevant context but were not re-labeled as BMP-related.

##### Amyotrophic lateral sclerosis (ALS)

The ALS analysis used Hop et al. as the primary dataset because it is the largest harmonized ALS exome rare-variant analysis discussed in this project and it explicitly reports geneset burden results in addition to gene-level findings. The paper analyzed 22 cohorts and reported URV geneset burden analyses across 11,777 GO, KEGG and Reactome gene sets from MSigDB v7.5, after excluding gene sets with fewer than 5 or more than 1,000 genes.

The pathway-level BMP audit examined the paper's own filtered geneset results after removal of single-gene-driven pathway signals. According to the study, only two genesets remained significant after this filtering step: "regulation of mRNA splicing via spliceosome" and its parent term "regulation of RNA splicing". No BMP-related term was reported among the significant or conditional pathway findings. The article text itself was additionally queried for the strings BMP and bone morphogenetic, which yielded no pathway-level evidence of BMP involvement.

The secondary gene-level audit intersected the fixed BMP catalog with the exome-wide significant and replicated genes emphasized by the authors, the genes contributing to the significant RNA-splicing signal, and the candidate-gene GO follow-up reported in the supplementary material. ALS was classified as not supportive of BMP overrepresentation because the pathway-level result was negative and the highlighted ALS genes did not reveal a canonical BMP cluster.

##### Parkinson's disease (PD)

The primary PD comparison used the multi-ancestry GWAS meta-analysis of Kim et al. because it is the largest PD common-variant dataset discussed here with explicit MAGMA/FUMA pathway analysis. The paper tested 16,992 MSigDB v7 gene sets, adjusted pathway results using Benjamini–Hochberg FDR, and then performed conditional analysis to remove redundant terms. This published pathway framework was treated as the decisive enrichment layer for PD.

The pathway-level BMP audit screened the corrected-significant ontology terms reported by Kim et al. for any explicit BMP-related label. The same paper highlighted significant non-BMP categories involving immune pathways, mitochondria, vesicles/phagolysosome-autophagosome biology and tau protein kinase activity, but no corrected BMP-specific pathway term was reported. The article text was additionally queried for BMP-related keywords and did not provide pathway-level evidence for BMP enrichment.

The secondary gene-level audit intersected the fixed BMP catalog with the 12 potentially novel loci named in the paper, the 6 loci fine-mapped to a single putative causal variant, and the 25 putative genes nominated from the novel loci by eQTL/SMR analyses. As an orthogonal PD-specific check, the same symbol-level overlap procedure was also applied to the highlighted findings from two large PD rare-variant studies: the familial exome analysis identifying RAB32 and the large replication meta-analysis dominated by GBA1 and LRRK2 signals. PD was classified as not supportive of BMP overrepresentation because neither the primary multi-ancestry pathway layer nor the secondary rare-variant checks yielded a BMP-centered signal.

#### Datasets

**A1. Alzheimer's disease gene set**

Derived from locus-prioritized genes in Bellenguez et al. 2022

APOE
BIN1
PICALM
CR1
CLU
ABCA7
TREM2
CD2AP
EPHA1
MS4A6A
CD33
SORL1
FERMT2
INPP5D
MEF2C

HLA-DRA
CASS4
PTK2B
SLC24A4
RIN3
ZCWPW1
NME8
CELF1
SPI1
ADAM10
IQCK
ACE
ADAMTS1
WWOX
ECHDC3
APP
SHARPIN
PLCG2
ABI3
DOC2A
ICA1
ICA1L
DGKQ
WDR81
LIME1
EGFR
TSPAN14
TMEM106B
GRN
TNIP1
NCK2
RBCK1
OTULIN
BLNK
SIGLEC11
RHOH
LILRB2
RASGEF1C
SPPL2A
HS3ST1
APH1B
UNC5CL
CNTNAP2
SORT1
CTSB
MME

PILRA
PILRB
SCIMP
NTN5
HAVCR2
JAZF1
CCDC6
MAF
GAB2

---

**A2. Amyotrophic lateral sclerosis gene set**

Derived from gene-level results in ALS exome study 2026

SOD1
TARDBP
FUS
TBK1
NEK1
KIF5A
OPTN
VCP
UBQLN2
DNAJC7
C9orf72
SQSTM1
CCNF
ANXA11
TIA1
CHCHD10
KIF4A
KNTC1
YKT6
HTR3C
GBGT1
ARPP21
CFAP410
CAPN2

---

**A3. Parkinson's disease gene set**

Derived from loci and fine-mapped genes in Kim et al. 2024

SNCA
LRRK2
GBA1
MAPT
TMEM175
GCH1
BST1
RAB29
VPS13C
DNAJC13
ATP13A2
FBXO7
PLA2G6
PINK1
PRKN
DJ1
TMEM163
HIP1R
LSM7
CAMK2D
MTF2
PIK3CA
ADD1
SYBU
IRS2
USP8
PIGL
FASN
MYLK2
USP25
EP300
PPP6R2
RAB32

#### How the negative conclusion was reached

The phrase "BMP genes were not overrepresented" was not used as a statement about absolute biological impossibility. Instead, it was used in the narrower and reproducible sense defined in this note: the major published human-genetics studies selected for AD, ALS and PD did not show BMP overrepresentation within their own multiple-testing-corrected pathway analyses, and the papers' highlighted disease-gene outputs did not reveal a convincing secondary concentration of BMP genes on exact symbol overlap audit.

The pathway layer carried the most weight in the final interpretation because it is the appropriate study-internal test for overrepresentation. This was especially important for AD and PD, where the source papers already performed MAGMA competitive pathway analysis, and for ALS, where the source paper already performed large-scale geneset burden analysis. The gene-overlap layer was used as a confirmatory check rather than as the primary inferential engine.

The comparison therefore asked a deliberately conservative question: do the best available disease-genetics studies themselves nominate BMP signaling as a corrected, reproducible enriched pathway? Because the answer was no in each case, and because the secondary gene-level overlap audit did not overturn that conclusion, the overall interpretation was that BMP overrepresentation was not supported by the selected AD, ALS and PD datasets.

#### Minimal reproducible screening algorithm (pseudocode)

The exact logic of the comparison can be summarized as follows:

```
bmp_catalog = fixed_209_gene_catalog()  # imported unchanged from companion file

for study in [AD_Bellenguez_2022, ALS_Hop_2026, PD_Kim_2024]:

    pathways = load_corrected_significant_pathways(study)

    bmp_pathway_hit = any(explicitly_bmp_related(term) for term in pathways)

    disease_genes = normalize_symbols(load_reported_disease_genes(study))

    bmp_gene_overlap = sorted(set(disease_genes) & set(bmp_catalog))

    if study == PD_Kim_2024:

        for secondary_pd_study in [PD_Hop_2024, PD_Pitz_2024]:

            disease_genes |=
normalize_symbols(load_reported_disease_genes(secondary_pd_study))

            bmp_gene_overlap = sorted(set(disease_genes) & set(bmp_catalog))

    call_negative = (bmp_pathway_hit is False) and (bmp_gene_overlap is not convincing)

    report(study, bmp_pathway_hit, bmp_gene_overlap, call_negative)
```

Here, “explicitly_bmp_related(term)” means that the term label itself is BMP-specific (for example, contains BMP or bone morphogenetic), not merely broadly developmental or TGF-β-related. “Not convincing” in the last line means that the gene-level overlap did not reveal a coherent BMP cluster capable of overturning the negative pathway-level result.

#### Complete Python script

Disease-associated gene sets were derived directly from the supplementary materials of the respective studies (Bellenguez et al. 2022; Kim et al. 2024; ALS exome study 2026) by extracting locus-prioritized or exome-significant genes as defined by the authors.”

```
import pandas as pd
import numpy as np
from scipy.stats import hypergeom
from statsmodels.stats.multitest import multipletests
# -----------------------------
# CONFIGURATION
# -----------------------------
# Universe size (protein-coding genes)
GENE_UNIVERSE_SIZE = 20000
# Input files (simple one-column gene symbol lists)
```

```
BMP_FILE = "bmp_genes.txt"
DATASETS = {
    "AD_Bellenguez_2022": {
        "genes_file": "ad_genes.txt",
        "bmp_pathway_detected": False
    },
    "ALS_exome_2026": {
        "genes_file": "als_genes.txt",
        "bmp_pathway_detected": False
    },
    "PD_Kim_2024": {
        "genes_file": "pd_genes.txt",
        "bmp_pathway_detected": False
    }
}
# -----------------------------
# FUNCTIONS
# -----------------------------
def load_gene_list(path):
    genes = pd.read_csv(path, header=None)[0]
    return set(genes.str.upper().str.strip())
def hypergeometric_test(bmp_genes, disease_genes, universe_size):
    """
    Test whether BMP genes are enriched in disease gene list.
    """
    M = universe_size
    n = len(bmp_genes)
    N = len(disease_genes)
    k = len(bmp_genes & disease_genes)
    # P(X >= k)
    pval = hypergeom.sf(k - 1, M, n, N)
    expected = (n / M) * N
    if expected > 0:
        fold_enrichment = k / expected
    else:
        fold_enrichment = np.nan
    return {
        "overlap_k": k,
        "expected": expected,
        "fold_enrichment": fold_enrichment,
        "p_value": pval
    }
# -----------------------------
# MAIN ANALYSIS
# -----------------------------
bmp_genes = load_gene_list(BMP_FILE)
results = []
for dataset_name, info in DATASETS.items():
    disease_genes = load_gene_list(info["genes_file"])
    stats = hypergeometric_test(
        bmp_genes,
        disease_genes,
        GENE_UNIVERSE_SIZE
    )
    results.append({
        "dataset": dataset_name,
        "n_disease_genes": len(disease_genes),
        "bmp_overlap": stats["overlap_k"],
        "expected_overlap": stats["expected"],
        "fold_enrichment": stats["fold_enrichment"],
        "p_value": stats["p_value"],
        "bmp_pathway_detected": info["bmp_pathway_detected"]
    })
df = pd.DataFrame(results)
# -----------------------------
# MULTIPLE TESTING CORRECTION
```

```
  # ----------------------------
  df["FDR"] = multipletests(df["p_value"], method="fdr_bh")[1]
  # ----------------------------
  # DECISION RULE
  # ----------------------------
  def call_enrichment(row):
      if row["bmp_pathway_detected"]:
          return "YES (pathway-level)"
      if row["FDR"] < 0.05 and row["fold_enrichment"] > 1.5:
          return "YES (gene-level)"
      return "NO"
  df["BMP_enrichment_call"] = df.apply(call_enrichment, axis=1)
  # ----------------------------
  # SAVE OUTPUT
  # ----------------------------
  df.to_csv("bmp_enrichment_results.csv", index=False)
  print("\n=== RESULTS ===\n")
print(df)
```
